# Cerebellar outputs for rapid directional refinement of forelimb movement

**DOI:** 10.1101/2025.10.01.679895

**Authors:** Ayesha R. Thanawalla, Oren Wilcox, Eliza Rhee, Juan Jiang, Kee Wui Huang, Ephraim Boamah, Raihana Yusufi, Dalia Saklaway, Mohamed ElTabbal, James M. Conner, Akira Nagamori, Albert I. Chen, Eiman Azim

## Abstract

The cerebellum is critical for movement coordination, yet how outputs from the cerebellum continually guide the limbs and whether discrete pathways differentially contribute to adjusting motor output remain unclear. Using intersectional viral approaches in mice, we identified two spatially intermingled yet anatomically distinct neuronal populations in the cerebellar nuclei that drive the forelimb either toward or away from the body. Neural recordings revealed cerebellar activity that correlates with and precedes opposing directional changes in limb movement. Both cerebellar output pathways influenced motor neuron and muscle activity within milliseconds, producing reliable effects on limb trajectory despite substantial underlying variability in muscle recruitment patterns. Our findings disentangle a subtype organization to cerebellar limb control, revealing a subcortical circuit basis for online directional refinement during movement execution.

---

Moving with dexterity under a wide variety of behavioral and environmental conditions relies on sensorimotor circuits that adjust motor output rapidly to account for movement variability (*1, 2*). The cerebellum is critical for this process of online refinement (*3–11*). To impact movement, the cerebellum must ultimately recruit downstream motor circuits that then engage appropriate combinations of muscles. The predominant output pathways through which the cerebellum can affect limb movements are located in the cerebellar nuclei (CN) (*12, 13*). Perturbation of CN neurons produces pronounced limb motor deficits (*12, 14–21*), and abnormal CN activity is sufficient to generate a variety of movement pathologies including ataxia, dystonia, and tremor (*22*). Electrophysiological studies have linked CN neuronal activity to limb kinematics and rapid responses to movement perturbations (*19, 23, 24*), suggesting a role in tuning ongoing movements. Yet, whether there is a functional logic to cerebellar outputs, with discrete pathways dedicated to distinct aspects of online control, remains unclear.

CN neurons send extensive axonal projections throughout the brain and spinal cord (*12, 13, 20, 25–29*). This daunting anatomical complexity, combined with the challenges of isolating specific pathways, has made it difficult to explore whether a modular functional organization for cerebellar movement control might exist (*12, 30, 31*). Cerebellar communication with the neocortex via the thalamus implicates these ascending cerebellar pathways in learning, preparation, and execution of movement (*16, 18, 21, 32–36*). Yet the impacts of other cerebellar output pathways on the limb remain far less explored. In mice, activation of inputs to the cerebellum can modulate forelimb motor neuron activity and disrupt movement at very short latencies (*37*), and silencing CN neurons with direct projections to the spinal cord affects performance in a forelimb reaching task (*20*). These findings suggest that CN output pathways caudal to the thalamus, with faster access to spinal motor neurons, can exert rapid effects on the forelimb. The organization and function of these circuits, however, remain unresolved.

## CN neurons are organized into distinct output networks

The CN can be subdivided by histological and anatomical criteria into the fastigial (medial in rodents), interposed, and dentate (lateral in rodents) nuclei, which can be further divided into subnuclei (*38*). We reasoned that if functional distinctions in cerebellar outputs from these nuclei exist, they might be revealed by examining pathways with disparate rostro-caudal projection targets and, therefore, distinct routes to motor neurons: ascending projections to the motor thalamus (ventral anterior lateral complex, VAL) and descending projections to the cervical spinal cord (cSC) (*12, 13, 20*). We performed dual retrograde viral tracing from the VAL and cSC (segments cervical C4 to thoracic T1) and labeled neurons were quantified across the lateral (Lat), interposed (Int), and medial (Med) CN (**Fig. 1A**). CN neurons projecting to VAL were abundant (mean = 1699 neurons per animal ± 207 SEM) and widespread across subnuclei, while CN neurons projecting to cSC were far fewer in number (mean = 162 neurons per animal ± 43 SEM) and were found predominantly in the subnuclei of the Med and Int (**Fig. 1A**). Consistent with previous work (*20*), we found CN neurons that project to both VAL and the cSC across several subnuclei, most predominantly in the Int and Med subnuclei (5.37 ± 1.52% of all VAL projecting CN neurons and 37.46 ± 7.16% of all cSC projecting neurons) (**Fig. 1A-C**). These findings highlight the VAL as a major recipient of cerebellar output, receiving convergent input from a large population of neurons spread across the CN. Moreover, we identified three subgroups within the CN: a) neurons that project to VAL but not the cervical spinal cord (**CN^VAL^**); b) neurons that project to both VAL and cervical spinal cord (**CN^VAL/cSC^**); and c) neurons that target the cervical spinal cord but not VAL (**CN^cSC^**).

**Fig. 1.**
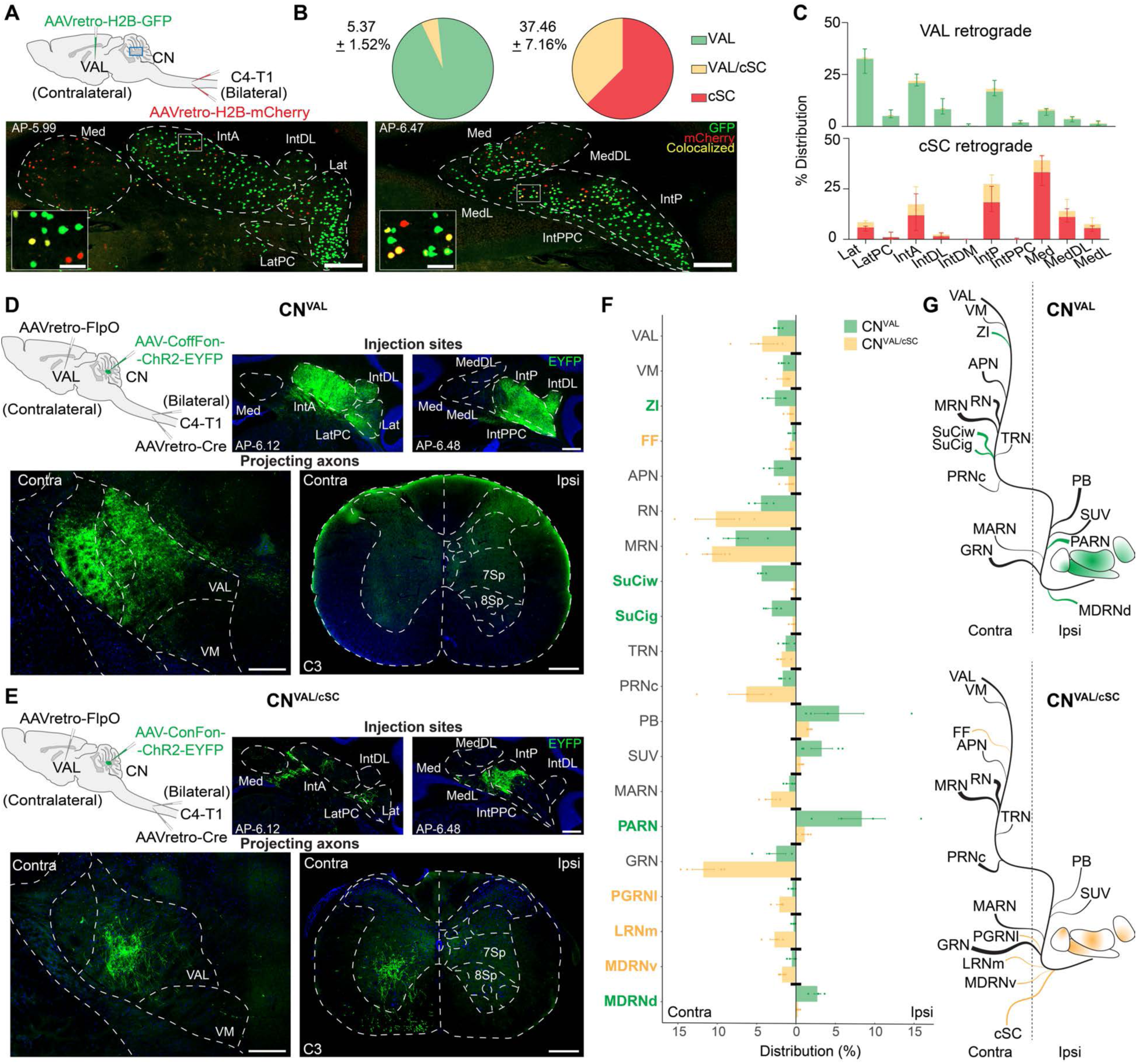
Retrograde targeting of VAL and cSC projecting CN neurons reveals distinct output networks. **(A)** Double retrograde mapping of VAL and cSC projecting neurons (top, blue box indicates CN region shown in images). Retrogradely labeled VAL projecting neurons (green; GFP+), cSC projecting neurons (red; mCherry+), and double-labeled VAL/cSC projecting neurons (yellow; GFP+/mCherry+) in the CN (bottom). Scale bars: 250 μm; inset, 50 μm. AP coordinates indicate distance from bregma. Lateral (Lat), lateral parvicellular (LatPC), interposed anterior (IntA), interposed dorsolateral (IntDL), interposed dorsomedial (IntDM), interposed posterior (IntP), interposed posterior parvicellular (IntPPC), medial (Med), medial dorsolateral (MedDL), lateral medial nucleus (MedL). **(B)** Percentage of double-labeled neurons (yellow) within the VAL projecting (left) and cSC projecting (right) populations (N=6, mean ± SEM). **(C)** Distribution of retrogradely labeled neurons across CN subnuclei. Double-labeled neurons (yellow) are represented as a fraction within each subnucleus (N=6, median and IQR). **(D)** Strategy for intersectional targeting of CN^VAL^ neurons (top, left). Expression of ChR2-EYFP in the CN at the injection sites (top, right). ChR2-EYFP expression in axons projecting to the VAL and lack of axons projecting to the cSC (bottom) in an example animal. Scale bars: 250 μm. **(E)** Strategy for intersectional targeting of CN^VAL/cSC^ neurons (top, left). Expression of ChR2-EYFP in the CN at the injection sites (top, right). ChR2-EYFP expression in axons projecting to the VAL and the cSC (bottom) in an example animal. **(F)** Distribution of axonal projections across brain regions, expressed as a percentage of total pixels detected within each animal. A subset of identified regions with high density projections consistent across animals within each group are represented here (Materials and Methods). Target regions highlighted in color are differentially present in one group versus the other (CN^VAL^, green, N=4; CN^VAL/cSC^, yellow, N=4; mean ± SEM). Brain region abbreviations and quantification are described in **table S1**. **(G)** Schematics depicting CN^VAL^ and CN^VAL/cSC^ cerebellar output pathways. The thickness of the lines represents relative density within each group. Lines indicated in color represent distinguishing projections of each pathway.

The existence of neurons that project to both thalamus and spinal cord (**Fig. 1A-C**) meant that we could not use a simple retrograde targeting strategy to access specific subgroups, as single injections into either VAL or cSC would target more than one of the subgroups we identified. We, therefore, used intersectional viral strategies (*39*) to target the three distinct populations in a Boolean manner with a membrane targeted fluorophore (ChR2-EYFP) (**Fig. 1D**,**E** and **fig. S1**), and we quantified labeled axons using the QUINT workflow for atlas registration and region-specific quantification of fluorescent signals (*40*). By measuring the distribution of projections within each mouse across all brain regions as well as projection density within each region, we found that both populations of neurons projecting to the VAL (CN^VAL^ and CN^VAL/cSC^) exhibit extensive collateralization, including to targets in the zona incerta, red nucleus, midbrain reticular region, pontine areas, and rostral and caudal medulla (**Fig. 1F,G, fig. S2**, and **table S1**). While many regions receive projections from both CN^VAL^ and CN^VAL/cSC^ neurons, we identified target areas where the two populations diverge, several of which have been implicated in forelimb movement (*41, 42*). CN^VAL^ neurons send unique projections to the superior colliculus and preferentially target the lateral column of the ipsilateral medulla (**Fig. 1F,G, fig. S2A,C,E, fig. S3**, and **table S1**), while CN^VAL/cSC^ neurons are distinguished by projections to the cSC and preferentially target the medial column of the contralateral pontine/medullary region (**Fig. 1F,G**, **fig. S2B,D,E, fig. S3**, and **table S1**). In contrast, the CN^cSC^ population sends bilateral projections to the intermediate laminae of the cervical spinal cord with fewer projections in the brain (**fig. S1**). Thus, intersectional anatomical targeting reveals subpopulations of CN neurons that target diverse premotor areas, many with direct projections to motor neurons in the cervical spinal cord (*37, 41, 43–46*), providing potential routes for rapid influence on forelimb movements.

## CN subpopulations drive opposing forelimb movements

To test whether functional distinctions exist between these anatomically divergent pathways, we trained mice in a head fixed water reaching behavioral task (*47*) (**Fig. 2A** and **Movie 1**) and used the intersectional viral approaches described above to selectively target each group with the excitatory opsin ChR2 (**Fig. 2B,C** left and **fig. S4A** left). Mice were trained to reach for a water droplet in response to an auditory cue, and each CN subpopulation was photostimulated on a pseudo-randomized 20–25% subset of reach trials (18 Hz, 5 ms pulse width, 473 nm). Trials began when the right hand was at rest on a metal perch sensor, which served as a trigger to initiate optogenetic perturbation when the hand was lifted at reach onset (Materials and Methods). Dependent on reach duration, each trial consisted of ∼4-5 stimulation pulses.

**Fig. 2.**
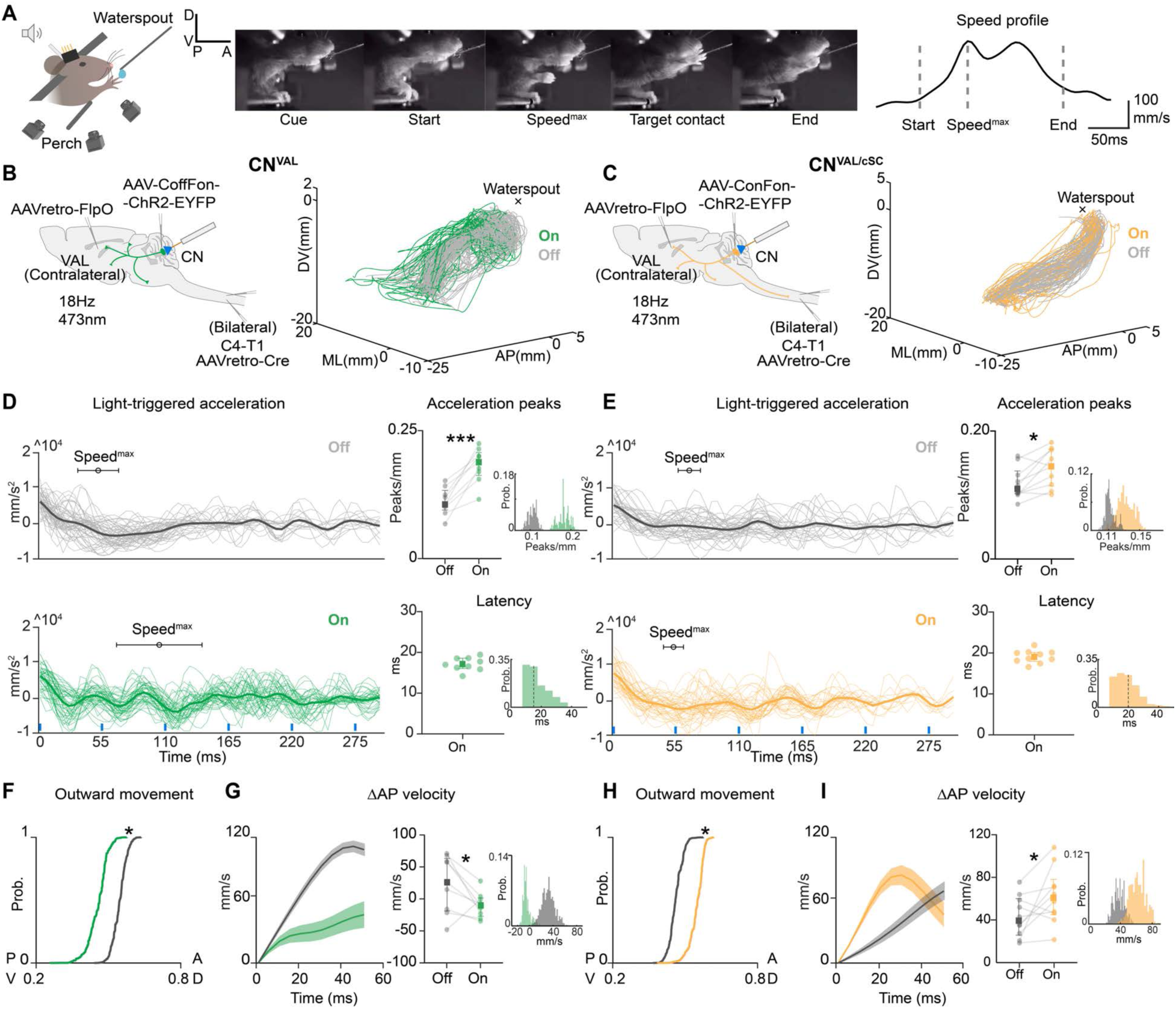
CN networks drive limb movements in opposing directions. **(A)** Schematic of head fixed water reaching setup (left), montage of an example reach (middle), and speed profile of an example reach with reach events marked: Start, Speed^max^, End (right). **(B)** Intersectional strategy to target CN^VAL^ neurons for expression of ChR2-EYFP (left). 3D trajectories of all reach trials from an example animal; grey lines: light off trials, green lines: light on trials (right). **(C)** Same as (B) for CN^VAL/cSC^ neurons; grey lines: light off trials, yellow lines: light on trials. **(D)** Hand acceleration triggered at light onset in CN^VAL^ trials (green) compared to similar timepoints during light off trials (grey) from a representative animal; thicker lines are mean, thinner lines are single trials, blue lines indicate pulses of light, distributions of when Speed^max^ occurred are shown as mean ± SEM (left). Number of acceleration peaks normalized to path length across animals (top right; N=9; p=0.0001, paired t-test); circles represent averages within animal, squares and error bars represent median and IQR across mice. Hierarchical bootstrapped distribution of medians (inset). Latency to acceleration onset (bottom right); circles represent trial averages within animal, error bars represent median and IQR across mice, distribution across mice (inset) with dotted line at median. **(E)** Same as (D) for CN^VAL/cSC^ trials. Number of acceleration peaks normalized to path length across animals (top right; N=10; p=0.0094, paired t-test). Bootstrapped distribution of medians (inset). Latency to acceleration onset (bottom right); distribution across mice (inset). **(F)** AP-DV component of the first light-triggered submovement (green) compared to the first reach submovement during light off trials (grey) across all CN^VAL^ animals (hierarchical bootstrapped CDF of medians; N=9; P-boot=0.0253, Off>On, by joint probability). **(G)** Change in AP velocity after light onset during light on trials compared to a similar timepoint during light off trials in a CN^VAL^ mouse (left). Lines indicate means, shaded regions indicate SEM. Change in AP velocity calculated in a 5 ms window 20–25 ms after light onset during light on trials compared to a similar timepoint during light off trials across CN^VAL^ mice (right; N=9; p=0.037, paired t-test); circles: averages within animal, squares and error bars: median and IQR across mice. Hierarchical bootstrapped distribution of medians (inset). **(H)** Same as (F) for CN^VAL/cSC^ animals (hierarchical bootstrapped CDF of medians; N=10; P-boot=0.0256, On>Off, by joint probability). **(I)** Same as (G) for a CN^VAL/cSC^ mouse. Change in AP velocity calculated in a 5 ms window 20–25 ms after light onset across CN^VAL/cSC^ mice (right; N=10; p=0.047, paired t-test). Hierarchical bootstrapped distribution of medians (inset).

CN^VAL^ neuronal stimulation resulted in retraction of the limb toward the body following each light pulse (**Movie 2**), while CN^VAL/cSC^ neuronal stimulation drove protraction away from the body (**Movie 3**). Stimulation of both CN^VAL^ and CN^VAL/cSC^ neurons produced abrupt changes in limb acceleration, generating deviations in reach trajectories when compared to light off trials (number of acceleration peaks/mm: CN^VAL^; light off: median = 0.106, interquartile range (IQR) = 0.038, light on: median = 0.189, IQR = 0.044 | CN^VAL/cSC^; light off: median = 0.110, IQR = 0.035, light on: median = 0.145, IQR = 0.058) (**Fig. 2B-E**). Moreover, these kinematic perturbations occurred rapidly after light onset (CN^VAL^; median = 16.98 ms, IQR = 2.52 | CN^VAL/cSC^; median = 19.16 ms, IQR = 2.2 ms) (**Fig. 2D**,**E** right). In contrast, we did not find comparable effects when stimulating CN^cSC^ neurons (CN^cSC^; light off: median = 0.118 peaks/mm, IQR = 0.027, light on: median = 0.109 peaks/mm, IQR = 0.055) (**Movie 4** and **fig. S4**) or in control animals with expression of a fluorophore in excitatory CN neurons (light off: median = 0.106 peaks/mm, IQR = 0.028, light on: median = 0.111 peaks/mm, IQR = 0.015) (**fig. S5**). One possibility is that CN^cSC^ neurons, which predominantly localize to the rostral Med nucleus, are involved in balance or postural control (*12*), which cannot be clearly resolved in a head fixed reaching assay.

Because activation of CN^VAL^ and CN^VAL/cSC^ neurons appeared to drive the forelimb in opposing directions, we next focused on these populations and quantified how limb movements are affected along the axes of the reaching behavior. We examined the segment of movement immediately following the first light stimulation until the next break in movement and computed the average unit displacement vector. Stimulation of CN^VAL^ neurons reduced outward displacement, when compared to the first submovement during unstimulated reaches (defined by local velocity minima, Materials and Methods) (**Fig. 2F**), while stimulation of CN^VAL/cSC^ neurons increased it (**Fig. 2H**) (anterior-posterior/dorsal-ventral (AP-DV) component of the vector: CN^VAL^; light off: median = 0.553, IQR = 0.043, light on: median = 0.472, IQR = 0.059 | CN^VAL/cSC^; light off: median = 0.459, IQR = 0.086, light on: median = 0.551, IQR = 0.041). To determine whether these CN populations have the same effect when the limb is moving toward the body, we identified timepoints when animals generated spontaneous inward movements and found that CN^VAL^ stimulation again pulled the limb toward the body while CN^VAL/cSC^ stimulation pushed the limb outward (**fig. S6**) (AP-DV component of the vector: CN^VAL^; light off: median = -0.006, IQR = 0.087, light on: median = -0.163, IQR = 0.082 | CN^VAL/cSC^; light off: median = -0.088, IQR = 0.029, light on: median = 0.060, IQR = 0.051). Thus, these CN output pathways drive specific directional adjustments of the limb toward or away from the body, irrespective of the ongoing direction of limb movement.

We next assessed how neuronal activation influenced outward limb velocity and found that CN^VAL^ stimulation reduced ensuing AP velocity (**Fig. 2G**), while CN^VAL/cSC^ stimulation increased it (**Fig. 2I**) (ΔAP velocity: CN^VAL^; light off: median = 26.240 mm/s, IQR = 82.230, light on: median = -9.836 mm/s, IQR = 28.452 | CN^VAL/cSC^; light off: median = 39.32 mm/s, IQR = 34.310, light on: median = 60.75 mm/s, IQR = 33.100). The kinematic effects were most consistent in the AP dimension, toward and away from the body (**Fig. 2G,I** and **fig. S7**), prompting us to compute the relative AP displacement of the forelimb throughout the entire reach, encompassing all light pulses. CN^VAL^ stimulation reduced overall anterior displacement, a consequence of inducing submovements that oppose the natural reach direction toward the target (AP displacement ratio : CN^VAL^; light off: median = 6.061, IQR = 1.767, light on: median = 2.965, IQR = 0.547) (**fig. S8A**). In contrast, stimulation of CN^VAL/cSC^ neurons did not disrupt total anterior displacement across the full duration of the reach, given that these neurons promote submovements in the same direction as the target (AP displacement ratio: CN^VAL/cSC^; light off: median = 4.432, IQR = 1.030, light on: median = 5.487, IQR = 1.844) (**fig. S8B**). Together, these data reveal antagonistic roles for two anatomically discrete populations of CN neurons that can rapidly drive limb movement toward or away from the body.

## CN neuronal activity correlates with bidirectional changes in forward movement

While targeted activation of two classes of CN neurons drives the limb in opposing directions, it remained unclear whether the physiological activity of CN neurons during natural reaching reflects similar functional distinctions. As has been seen for limb deceleration (*19*), we hypothesized that during reaching we would be able to detect subpopulations of CN neurons whose activity correlates with and precedes rapid adjustments to forelimb movement in both the anterior and posterior directions. We thus performed electrophysiological recordings with tetrodes (8 bundles, 32 channels per probe) chronically implanted in the Int and Med nuclei of mice performing head fixed water reaching (**fig. S9A-C**), finding units whose activity was consistently modulated across reaches (218 units across 6 mice) (**Fig. 3A**).

**Fig. 3.**
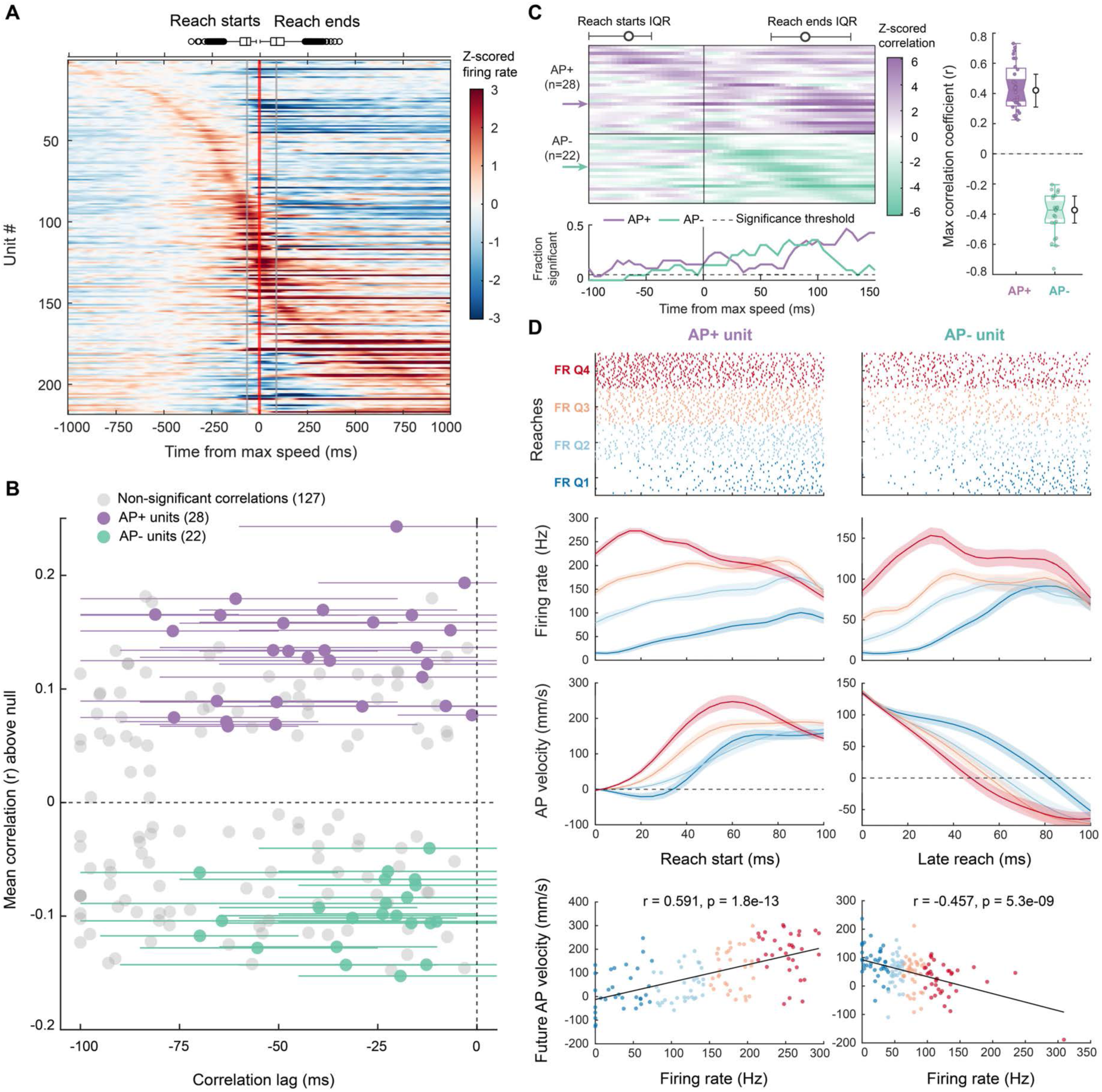
CN neuronal activity correlates with bidirectional changes in forward limb movement. **(A)** CN units modulate their activity during reaches (z-scored firing rates shown). Reach starts and ends are indicated (line, median; boxes, IQR; whiskers, non-outlier minima & maxima; open circles, outlier minima & maxima) relative to maximum hand speed at time zero. All sessions and animals (N=6 mice, 218 units). **(B)** Positive (top) and negative (bottom) correlations with AP velocity above the trial shuffled null of all units across time lags. Units with positive correlations (AP+, purple, N=28), negative correlations (AP-, teal, N=22), and non-significant correlations (grey), with whiskers on AP+ and AP-units indicating the lag range over which the correlation was significant. **(C)** Sliding window z-scored correlations between firing rate and future AP velocity relative to maximum hand speed for all AP+ and AP-units, with median and IQR of reach starts and ends (top). Each unit’s correlation lag was determined from (B). Arrows indicate units shown in (D). Fraction of significantly correlated AP+ and AP-units over time, relative to maximum hand speed (bottom). Maximum and minimum correlation coefficients for AP+ and AP-units with hierarchical bootstrapped median values and 95% confidence intervals (right). **(D)** Example unit rasters across all reaches binned into firing rate (FR) quartiles (defined by 0 to +50 ms after time zero, top), smoothed firing rate (middle top), AP velocity (middle bottom). Left shows an example AP+ unit relative to reach start and right shows an example AP-unit relative to a time point in late reach when AP velocity drops below 130 mm/s. Bottom plots show AP+ and AP-unit correlations of FR with future AP velocity, determined by each unit’s cross-correlation lag in (B), fit with linear models (r and p values shown). Also see **fig. S9H-J**.

To examine the relationship of unit activity to forelimb kinematics, we performed cross-correlation of the estimated instantaneous firing rate with the AP velocity of the hand across all acceleration and deceleration epochs during every reach, finding units with both positive and negative cross-correlation peaks at a broad range of time lags (**fig. S9D**). We then selected units whose activity was correlated or anticorrelated with forward velocity (referred to as AP+ and AP-units, respectively) at a negative time lag, where changes in neural activity are correlated with future changes in kinematics (**Fig. 3B**). To complement our cross-correlation, which captures linear relationships, we used a mutual information estimator (*48*) to quantify all dependencies between firing rate and AP velocity. The time lag at which mutual information between firing rate and AP velocity peaked was similar when compared to our cross-correlation analysis (**fig. S9E,F**), indicating that our linear approach effectively captured a major component of the relationship between firing rate and kinematics.

Based on our cross-correlation criteria, we next asked when during the reach did unit spiking predict changes in AP velocity. We performed a sliding window correlation between binned spike count and AP velocity on AP+ and AP-units at each unit’s lag, as identified in our cross-correlations. To compare units from different sessions, we z-scored the correlations against a trial-shuffled null correlation of the same data and quantified the fraction of correlations exceeding a significance threshold (Materials and

Methods). We found AP+ unit correlations that cover many phases of the behavior, with correlations for both populations appearing during the second half of the reach, when corrective submovements are often used to reduce residual error as the hand approaches the target (**Fig. 3C**) (*24, 49, 50*). Uncorrelated units showed no movement related patterns (**fig. S9G**). AP+ and AP-unit correlations were further confirmed by dividing the activity of units across reaches into firing rate quartiles at various phases of the reach to visualize how the firing rate corresponds to future kinematic changes (**Fig. 3D** and **fig. S9H-J**). We attempted to determine whether there is a link between the anatomically defined CN^VAL^ and CN^VAL/cSC^ groups and these AP+ and AP-classes through optotagging experiments, but we found that the efficiency of the intersectional viral approach limited yield (Supplementary Text). These findings demonstrate that subpopulations of CN neurons show changes in activity that correlate with and precede bidirectional changes in limb kinematics at several distinct phases of the reach, potentially providing a basis for online adjustments throughout movement execution.

## CN subpopulations flexibly and rapidly recruit forelimb muscles to drive movement

In principle, CN^VAL^ and CN^VAL/cSC^ output pathways could direct limb movement by invariably recruiting specific sets of muscles. One simple hypothesis for rigid muscle recruitment is that CN^VAL^ neurons, which pull the limb inwards, would activate flexors, while CN^VAL/cSC^ neurons, which propel the limb outward, would activate extensors. Alternatively, they could operate in a more flexible task-oriented space, driving different combinations of muscle activity to achieve similar kinematic output.

We performed EMG recordings from elbow and shoulder flexor (biceps brachii, anterior deltoid) and extensor (triceps brachii, posterior deltoid) muscles that are active during outward reaching (**Fig. 4A** and **fig. S10A**). When we examined normal EMG activity during unstimulated control trials, we found that the spatial (which muscle is recruited) and temporal (when the muscle is recruited) profiles varied widely (**fig. S10A)**. To determine whether kinematically similar reaches can be produced by disparate patterns of muscle activity, we calculated pairwise root mean square error (RMSE) over time-normalized reach trajectories to identify the most kinematically similar pairs within each animal (RMSE < 10^th^ percentile) (**fig. S10B**). Using these similar reaches, we then examined pairwise cosine distances between EMG patterns (higher cosine distance means greater dissimilarity) and found a wide range of muscle activity (cosine distance, coefficient of variation: median = 0.43, IQR = 0.21) (**fig. S10B-D**), confirming that even in this restricted head fixed reaching task, mice engage a variety of muscle activity patterns to drive consistent kinematic output. As expected, when comparing kinematically dissimilar trial pairs (RMSE > 90^th^ percentile), we found larger EMG cosine distances (**fig. S10C,E**), confirming that activity in the forelimb muscles we recorded is associated with movement kinematics. We next asked whether activation of CN^VAL^ and CN^VAL/SC^ pathways recruits fixed or flexible patterns of muscle activity. Comparing highly similar light-triggered kinematics within animals, as above, we found highly disparate EMG patterns (cosine distance, coefficient of variation: CN^VAL^; median = 0.67, IQR = 0.33 | CN^VAL/cSC^; median = 0.64, IQR = 0.31) (**Fig. 4B-D** and **fig. S11**), demonstrating that these CN output pathways flexibly recruit different combinations of muscles to drive consistent kinematic outcomes.

**Fig. 4.**
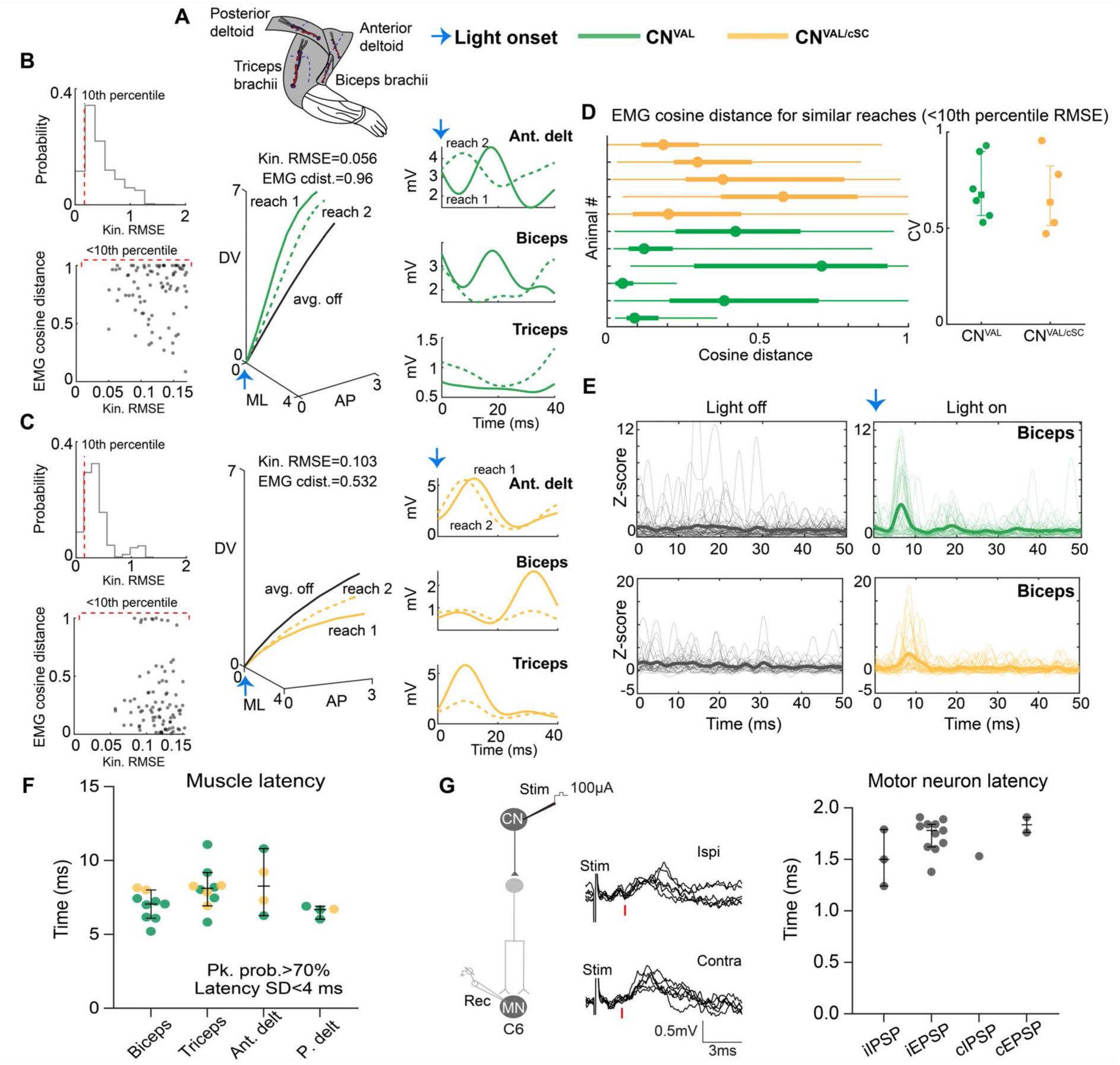
CN subpopulations flexibly and rapidly recruit forelimb muscles to drive bidirectional movements. **(A)** Schematic of forelimb muscles recorded during head fixed water reaching. **(B)** Distribution of RMSE values of pairwise kinematic (kin) comparisons in an example CN^VAL^ animal after light onset. Dotted red line indicates the most similar 10^th^ percentile cut-off value (top, left). EMG cosine distance (cdist) between reach pairs with RMSE values < 10^th^ percentile cut off (bottom, left). Example reach pair with low kinematic RMSE; reach 1, solid line; reach 2, dotted line; average light off trajectory (black) shown for reference (middle); scales in mm. EMG envelope for the two reaches (right). Light onset indicated by blue arrow. (Also see **fig**. **S11A**). **(C)** Same as (B) for an example CN^VAL-SC^ animal. (Also see **fig. S11A**). **(D)** Range of EMG cosine distance values for the most similar reach pairs (RMSE<10^th^ percentile cut-off). Each row is one animal, circle is median, thin line is range, thicker bar is IQR (left). Coefficient of variation (CV) of cosine distance values across animals (right). **(E)** Light-triggered averaging of z-scored EMGs from example animals in light off (left) and light on (right) conditions. Green, example CN^VAL^ mouse; yellow, example CN^VAL/cSC^ mouse (also see **fig. S12**). Thicker lines represent mean, thinner lines represent single trials. Light onset (blue arrow) aligned to zero in plots on right. **(F)** Latency to the first peak in z-scored EMGs after light onset (median and IQR across mice). Peak probability >70% within 20 ms of light onset and standard deviation of the latency of identified peaks < 4 ms were used as criteria (also see **fig. S12** and **table S2**). **(G)** Intracellular motor neuron (MN) recordings during electrical stimulation of the CN reveal modulation at disynaptic latencies. Example traces from ipsilateral (left, top) and contralateral (left, bottom) evoked EPSPs; each trace represents a single stimulation; red lines indicate EPSP onset. Latencies to fastest evoked EPSPs and IPSPs in ipsilateral (i) and contralateral (c) MNs (right; N=5 mice; median and IQR across MNs).

Ensuring accuracy during a fast movement like reaching implies an ability for cerebellar outputs to impact muscle activity rapidly. To quantify the latency of muscle recruitment, we compared event-triggered averaging of EMG signals after light onset to an equivalent point in the reach trajectory in unstimulated trials (**Fig. 4E** and **fig. S12**). Muscles recruited at the shortest latency were identified by setting strict thresholds for both the probability of muscle activation and variance in timing of muscle activity peaks after stimulation (**fig. S12A,B, table S2**, and Materials and Methods). With these criteria, we found rapid muscle recruitment when activating both CN subpopulations ranging from approximately 5 – 10 ms after light onset across all recorded muscles (biceps, median = 7.05 ms, IQR = 1.58; triceps, median = 8.12 ms, IQR = 1.19; ant. deltoid, median = 8.23 ms, IQR = 3.89; post. deltoid, median = 6.68 ms, IQR= 0.66 (**Fig. 4F**). These findings indicate that CN^VAL^ and CN^VAL/cSC^ output pathways are each capable of recruiting both elbow and shoulder flexors and extensors on very rapid timescales. To confirm these fast effects on muscles, we next determined whether CN neurons can recruit spinal motor neurons at such short latencies by performing intracellular recording from motor neurons in the cSC (C6) of anesthetized mice while electrically stimulating the Int nucleus. Stimulation evoked excitatory postsynaptic potentials (EPSPs) in the disynaptic range in 47.8% of ipsilateral motor neurons (median = 1.78 ms, IQR = 0.22; N = 23 neurons) and 13.3% of contralateral motor neurons (median = 1.84 ms, IQR = 0.15; N = 15 neurons) (**Fig. 4G**).

## Discussion

Through an intersectional anatomical approach, we identified distinct populations of cerebellar output neurons distinguished by their extensive collateral projections to subcortical premotor targets. Selective stimulation revealed a functional divergence of two CN subpopulations that drive rapid, opposing adjustments of limb movement toward or away from the body. These cerebellar outputs influence motor neurons and muscles within milliseconds, producing consistent kinematic outcomes through the flexible recruitment of muscles across joints. Complementary electrophysiological data support the idea that CN neurons parcellate into subpopulations whose activity correlates with and precedes bidirectional changes in limb kinematics. Collectively, these findings define cerebellar circuits for rapid directional control, providing neural substrates for the continuous online refinement of limb movement.

We initiated this study with the hypothesis that distinct functions for cerebellar outputs might be resolved by comparing populations that project to anatomically disparate downstream targets. This approach revealed subpopulations of CN neurons with diverse and broadly collateralizing projections to regions throughout the nervous system, providing further insight into the cell-type organization of cerebellar output (*12, 13, 20, 25, 26, 29*). Notably, these subpopulations are partially intermingled within the CN and do not strictly adhere to nuclear boundaries, highlighting the utility of pathway-specific targeting enabled by intersectional tools. While several of the downstream targets of the CN^VAL^ and CN^VAL/cSC^ pathways are shared, we identified sharp distinctions in some projection targets, for example in the superior colliculus and various reticular nuclei. It is likely that we uncovered just one of many anatomical branch points that demarcate CN subpopulations, and additional CN subtypes can be identified by similar intersectional strategies that leverage some of these other downstream structures as access points. Moreover, anatomical delineation of cerebellar output can continue to be informed by efforts to define the molecular heterogeneity of the CN throughout development and adulthood (*26, 51*). A question that cannot be answered with our approach is whether individual CN^VAL^ and CN^VAL/cSC^ neurons project collaterals broadly, or whether the diversity of targets we found is because the viral approach labeled populations of neurons, each of which might have more restricted projection profiles. Advances in tissue processing and microscopy that enable the reconstruction of axonal projections from individual neurons (*52*) can help to answer this question and provide insight into the anatomical and functional granularity of cerebellar outputs.

Thalamocortical pathways that receive input from the cerebellum have been implicated in diverse aspects of movement (*33–36, 53*). However, this ascending route cannot explain the disynaptic influence on motor neurons that we observed. We found that while both CN^VAL^ and CN^VAL/cSC^ neurons are defined by their projections to the thalamus, they also target brainstem and spinal regions that directly innervate motor neurons (*37, 41, 43, 45, 46*). Reticulospinal populations that are differentially innervated by CN^VAL^ and CN^VAL/cSC^ pathways represent compelling candidates (amongst several others in our anatomical data) for eliciting rapid and opposing effects on limb movement. Supporting this idea, we found that CN^VAL^ and CN^VAL/cSC^ neurons can be distinguished by their projections to MDRNv and MDRNd in the medulla, regions that have been implicated in driving either outward reach or inward hand-to-mouth movements (*41*). It is possible that the thalamic collaterals of CN^VAL^ and CN^VAL/cSC^ neurons also impact limb movement at longer latencies, or alternatively, these ascending projections could provide the neocortex with efference copies of the rapid motor commands emerging subcortically. We also note that the apparent numerical disparity between CN^VAL^ and CN^VAL/cSC^ neurons may be due to the fact that thalamus-projecting neurons are functionally diverse (*33, 35, 53*), with many neurons, especially in the Lat nucleus, involved in functions unrelated to limb movement. In contrast, CN^VAL/cSC^ neurons are defined by the much more stringent requirement that an individual neuron projects to thalamus and cervical spinal cord, revealing a smaller population located largely in the Int and Med nuclei that could be more functionally dedicated to limb control (*12*).

While our results show clear effects in the forward axis of the reaching movement, the logic of opposing subpopulations of CN neurons pushing in different directions is likely to apply more broadly. Behavioral tasks designed to promote forelimb movements with greater kinematic range may help to reveal a general framework for CN control in three dimensions. Moreover, directional control of movement by these cerebellar outputs could apply to other parts of the body. Discrete CN outputs controlling eyelid kinematics during closing and opening have been identified (*54*), and cerebellar cortical circuits upstream of the CN have been linked to controlling deceleration of the eye (*55*) and the tongue (*56*). It is, therefore, likely that delineating CN output circuits that affect the limb will provide insight into the online control of effectors across the body (*57*) and help to bridge the diverse movement pathologies related to cerebellar damage and disease with specific circuit dysfunction.

Our findings reveal that, even though CN neurons can recruit muscles within milliseconds, the spatial and temporal patterns of muscle recruitment vary. Moreover, the CN populations we identified can recruit both flexors and extensors depending on movement context—for example, during CN^VAL^ neuronal stimulation, both the biceps (elbow flexor) and posterior deltoid (shoulder extensor) can be activated as the limb retracts inward. Yet, despite variability in which muscles are recruited by CN neurons at any given time, the effects on how the limb moves toward or away from the body are consistent. These results suggest that the cerebellum can drive directional refinements in kinematic space through adaptable muscle activation, rather than through fixed recruitment of specific flexor or extensor muscles (*58, 59*). An interesting question for future study is how this phenomenon might apply to more innate behaviors like locomotion. Together, our findings suggest that evolutionarily ancient pathways connecting cerebellum, brainstem, and spinal cord are capable of using biomechanical degeneracy in the motor system (i.e., the capacity for many different combinations of muscle activity to drive similar task-level output) for flexible and rapid multi-joint control (*60–64*). Neocortical circuits that evolved more recently are likely to have leveraged the significant capacities of these subcortical circuits to enable and augment the online control of dexterous movement.

## Materials and Methods

### Mice

Procedures performed on mice were conducted according to US National Institutes of Health guidelines for animal research and were approved by the Institutional Animal Care and Use Committee of The Salk Institute for Biological Studies. Approximately equal numbers of adult (aged 2-6 months) male and female mice were used for all experiments; data were combined since no sex-specific differences were observed. All mice were maintained on a C57BL/6 background and housed on a 12:12 hour reverse light cycle. The following mouse lines were used: Wild-type (The Jackson Laboratory and in-house colony); *VGluT2-Cre* (B6J.129S6(FVB)-*Slc17a6*^tm2(cre)Lowl^/MwarJ; The Jackson Laboratory, 028863); R26-LNL-GtACR1 (R26-LNL-GtACR1-Fred-Kv2.1; B6;129S6 *Gt(ROSA)26Sor^tm3Ksvo^*/J, The Jackson Laboratory, 033089).

### Viruses

The following adeno-associated viruses (AAVs) were used, with serotype and titer (vg/ml) indicated: scAAVretro-hSyn-H2B-GFP, (Salk GT3 Core, 1.16 x 10^12^; plasmid: pAAV-hSyn-H2B-GFP, Callaway lab, Salk Institute for Biological Studies); scAAVretro-hSyn-H2B-mCherry, (Salk GT3 Core, 1.32 x 10^12^; plasmid: pAAV-hSyn-H2B-mCherry, Callaway lab, Salk Institute for Biological Studies); AAVretro-pmSyn1-Cre-EBFP (Salk GT3 Core, 1 x 10^12^; plasmid: AAV pmSyn1-EBFP-Cre, Addgene #51507, RRID: Addgene_51507); AAVretro-Ef1a-FlpO (Salk GT3 Core, 5 x 10^12^; plasmid: pAAV-EF1a-Flpo, Addgene #55637, RRID:Addgene_55637); AAV5-nEF-CoffFon-hChR2(H134R)-EYFP (Salk GT3 Core; 3 x 10^12^; plasmid: pAAV-hnEF Coff/Fon hChR2(H134R)-EYFP, Addgene #55649, RRID: Addgene_55649); AAV5-hSyn-Con/Fon-hChR2(H134R)-EYFP (Salk GT3 Core, 3 x 10^12^; plasmid: pAAV-hSyn Con/Fon hChR2(H134R)-EYFP, Addgene #55645, RRID: Addgene_55645); AAV5-EF1a-fDIO-EYFP (Salk GT3 Core, 2.6 x 10^12^; plasmid: pAAV-Ef1a-fDIO EYFP, Addgene #55641, RRID:Addgene_55641); AAV1-CAG-FLEX-tdtomato-WPRE (Penn Vector Core, 3 x 10^12^; plasmid: AAV pCAG-FLEX-tdTomato-WPRE, Addgene #51503, RRID:Addgene_51503); AAV8-EF1a-DIO-mCherry (Salk GT3 Core, 1.53 x 10^12^ ; plasmid: pAAV-EF1a-DIO-mCherry, Addgene #50462, RRID:Addgene_50462); AAV1-phSyn-flex-mRuby2-SypEGFP WPRE (Vigene Biosciences, 1.2 x10^12^; plasmid: AAV phSyn1(S)-FLEX-tdTomato-T2A-SypEGFP-WPRE, Addgene #51509, RRID:Addgene_51509).

### Antibodies

The following primary antibodies were used: rabbit anti-GFP (used for EGFP, EYFP; 1:1000; Thermo Fisher Scientific, A-11122); goat anti-GFP (used for EGFP, EYFP; 1:1000; Abcam, ab6673); rabbit anti-DsRed (used for mCherry, tdTomato; 1:1000; Living Colors, Takara Bio, 632496). The following conjugated secondary antibodies were used at a concentration of 1:500: donkey anti-goat-488 (Jackson Immuno Research Laboratories, 705-545-147); donkey anti-rabbit-488 (Jackson Immuno Research Laboratories, 711-545-152); donkey anti-rabbit-555 (Thermo Fisher Scientific, A-31572); biotin-SP donkey anti-goat (Jackson Immuno Research Laboratories, 705-065-147); biotin-SP donkey anti-rabbit (Jackson Immuno Research Laboratories, 711-065-152). For tyramide signal amplification (TSA), VECTASTAIN® Elite® ABC-HRP Kit, Peroxidase (Standard, Vector Laboratories, PK-6100) was used to make avidin-biotin-peroxidase complexes, and tyramide-conjugate Alexa Fluor™ 488 (1:1000; Thermo Fisher Scientific, B40953) was used as the substrate. NeuroTrace 640/660 (1:300; Thermo Fisher Scientific, N21483) was used as a Nissl stain.

### Electrode fabrication

*Intramuscular electromyography (EMG) electrode fabrication:* Bipolar EMG electrodes were custom made using Teflon coated 7 stranded stainless-steel wire (A-M systems, #793200) according to previously established protocols (*65, 66*). Approximately 25 cm of wire was looped around to create a double-stranded wire, and a knot was made 4 cm from one end of the wire strands; this length was determined based on the distance from the skull to the muscle of interest. On the longer side, 1 mm away from the knot, a small, bared region (recording site) ∼1 mm in length was made on each wire; the wires were then carefully twisted by hand, ensuring that the two recording sites were ∼1 mm apart. On the shorter side of the knot, each wire of an electrode pair was soldered to a pin of an 8-or 12-pin connector (Digikey, #SAM1161-04-ND). The two wire ends on the other side of the knot were crimped into the barrel of a 27- or 30-gauge needle, which was used for insertion into the muscle. The base of the connector with exposed wires was covered in epoxy (Gorilla 5-minute epoxy) to insulate and support the base of the connector. EMG electrodes were fabricated in either a four- or six-channel configuration.

*Tetrode drive fabrication:* Chronically implanted tetrode drives were fabricated using an adapted in-house protocol (*67*). Custom pieces were 3D printed on a Form 3 printer in Grey Pro or Grey V4 resin, washed in isopropanol, and UV cured. Stainless steel tubing of two diameters (18, 21 gauge) was inserted into the printed drive base and secured with epoxy (Gorilla 5-minute epoxy), and a printed hole in the drive base was threaded to hold a drive screw (18-8 Stainless Steel Narrow Cheese Head Slotted Screws, M1 x 0.25mm Thread, 10mm Long, McMaster-Carr, #91800A062). This assembly consisted of a base on which the drive body could smoothly travel vertically by turning a drive screw. An optical fiber (Fiber Optic Cannula, Ø1.25 x 6.4 mm Ceramic Ferrule, Ø200 µm Core, 0.39 NA, L=5 mm, Thorlabs, #CFMLC12L05) and eight 2 cm silicon tube segments (Molex Amber Silica, Polyimide Coated Smooth Solid Tubing, Capillary 0.004” (0.10 mm) 32.81’ (10.00 m), Digikey, #WM19371-ND) were inserted into the cannula on the drive body, such that the tubing and fiber protruded from the bottom of the drive assembly, and these components were fixed via epoxy to the moveable drive body. An EIB (electrode interface board, 36 pin Omnetics headstage, Neuralynx, #EIB-36-PTB) was mounted on two arms protruding from the drive base and secured via epoxy. Tetrode bundles were fabricated by twisting two 10 cm loops of tungsten wire (Tungsten 99.95% CS Wire, California Fine Wire Company, #100211) together using a custom Arduino spinner. Eight tetrode bundles were inserted into the eight silicone tubing segments until they protruded from the underside of the drive base alongside the optical fiber. Then, the wires in each tetrode bundle were inserted into the contact holes on the EIB and fixed in place with gold pins (Neuralynx, #31-0603-0100). The ventral protruding bundles were secured to the optical fiber with epoxy and then clipped to measure 0.5 mm longer than the tip of the optical fiber. The drive was then inserted into gold solution (Gold Plating Solution 10 ml/20 ml, Neuralynx, #GPS-10/20), connected to a NanoZ device (NanoZ Kit with holder, Neuralynx), and each wire was plated to a resistance of 70-90 kΩ. Next, a silver wire (PFA-Coated Silver Wire, 32/30 AWG, 25 ft, A-M Systems, #786500) was inserted into the reference and ground contact holes and pinned to both. One piece of 3D-printed shielding (to protect the implanted drive from damage) was cemented to the posterior end of the drive base.

### Surgical procedures

*General:* Stereotaxic surgical procedures performed on adult mice were carried out under isoflurane anesthesia (1.5%) on a precision stereotaxic frame (Kopf, Model 1900 Stereotaxic Alignment System). The surgical site was shaved and cleaned with betadine, alcohol, and sterile saline. Throughout all surgical procedures, animals were kept on a heating pad to maintain body temperature and eye lubricant was applied to protect the eyes from drying out during the procedure. Viral injections were performed using pulled glass capillaries and a Nanoject III (Drummond Scientific) mounted to a stereotaxic manipulator. For cerebellar nuclei and VAL thalamus injections, animals were placed in a stereotaxic frame in a flat skull position, and the skull was made level along the antero-posterior (AP) axis at bregma and lambda and medio-laterally (ML) ∼1 mm caudal from bregma. AP and ML coordinates were determined relative to bregma, and dorso-ventral (DV) coordinates were determined relative to the surface of the brain. A small craniotomy was made with a dental drill in the skull overlying the injection targets, and the tip of the glass capillary was lowered slowly into the brain.

For cervical spinal cord injections, animals were placed in the flat skull position, and the spine was stabilized by securing the tail of the animal to the stereotaxic base plate using tape to produce slight tension. An incision was made in the skin and superficial layers of muscle over the cervical spinal cord, and the skin and overlying muscles were then retracted, exposing the vertebrae. The muscle was cleaned from the surface of the spinal cord using a delicate bone scraper and fine cotton swabs. Injections were performed by piercing the dura with a 30-gauge syringe to facilitate penetration of the glass capillary into the spinal cord tissue. Injections were performed in between cervical vertebrae from above C4 to below C8, on either side of the central blood vessel, and DV coordinates were measured relative to the surface of the spinal cord. All viruses were injected at a rate of 3 nl/sec. Following all surgical procedures, overlying muscle layers and skin were sutured (6-0 black braided silk, Ethicon / 6-0 absorbable suture, Covetrus). Animals were given a subcutaneous injection of buprenorphine-ER (1 mg/kg) and Carprofen (5mg/kg) and were allowed to recover on a heating pad.

*Dual retrograde targeting of VAL and cSC projecting CN neurons to localize cell bodies:* To retrogradely label CN neurons, scAAVretro-hSyn-H2B-GFP or scAAVretro-hSyn-H2B-mCherry virus at the titers mentioned above were injected into either the VAL thalamus or cervical spinal cord. *VAL injections:* To cover the AP, ML, and DV extent of the VAL, virus was injected at 6 locations and at 4 DV positions. Coordinates for injection with respect to Bregma were as follows (all in mm): AP: -0.9, - 1.3, 1.7; ML (injected on left side): 0.9, 1.27. At each location, DV coordinates with respect to the surface of the brain were as follows: -3.8, -3.6, -3.4, - 3.2. The capillary was lowered to the most ventral location first, 20 nl of virus was injected, and the capillary was left in place for ∼20 s to prevent leak before moving dorsally to the next depth. *Cervical spinal cord injections:* To cover the AP, ML, and DV extent of cervical segments C4 to T1, virus was injected at 10 locations and at 5 DV positions. Virus was injected bilaterally at the midpoint between the central blood vessel and lateral extent of the cervical cord above C4, in between C4-C5, C5-C6, C6-C7, C7-C8, and below C8. At each location, DV coordinates with respect to the surface of the cord were as follows: -1.5, -1.3, -1.1, -0.9, -0.7. At each location the capillary was lowered to the most ventral location first, 25 nl of virus was injected, and the capillary was left in place for ∼20 s to prevent leak before moving dorsally to the next depth. To prevent any bias in labeling due to differences in viral efficiency, scAAVretro-hSyn-H2B-GFP and scAAVretro-hSyn-H2B-mCherry injections were randomized to the VAL or cervical spinal cord in each experimental animal.

### Intersectional targeting of CN networks

a. *CN^VAL^:* To specifically target *CN^VAL^* neurons, AAVretro-Ef1a-FlpO was injected into VAL thalamus, AAV5-nEF-CoffFon-hChR2(H134R)-EYFP was injected into the CN, and AAVretro-pmSyn1-Cre-EBFP was injected into the cervical spinal cord. This combination enabled the selective, conditional expression of ChR2-EYFP in neurons that project to the VAL, but not in those that project to the cervical spinal cord. To cover the extent of the VAL and segments C4-T1, viruses were injected at the locations and depths described above. To cover the extent of the cerebellar nuclei and subnuclei, 4 sites at two DV positions targeted the interposed anterior (IntA), interposed posterior (IntP), lateral (Lat), and medial (Med) nuclei. Coordinates for injecting the CN with respect to lambda were as follows: IntA: AP: -1.9, ML (injected on the right side): 1.44, DV: -2.5, -2.48; IntP: AP: -2.2, ML (injected right): 1.44, DV: - 2.5, -2.48; Lat: AP: -1.9, ML (injected right): 2, DV: -2.5, -2.48; Med: AP: -2.2, ML (injected right): 1, DV: -2.5, -2.48. At each location, the capillary was lowered to the most ventral location first, 40 nl of virus was injected, and the capillary was left in place for ∼ 30s to prevent leak before moving dorsally to the next depth.
b. *CN^VAL/cSC^:* To specifically target *CN^VAL/cSC^* neurons, AAVretro-Ef1a-FlpO was injected into the VAL, AAV5-hSyn-Con/Fon-hChR2(H134R)-EYFP was injected into the CN, and AAVretro-pmSyn1-Cre-EBFP was injected into the cervical spinal cord. This combination allowed selective conditional expression of ChR2-EYFP in neurons that project to both the cervical spinal cord and the VAL. To cover the extent of the VAL, segments C4-T1, and the CN, viruses were injected at locations and depths described above.
c. *CN^cSC^:* To specifically target CN^cSC^ neurons, AAVretro-Ef1a-FlpO was injected into the cervical spinal cord, AAV5-nEF-CoffFon-hChR2(H134R)-EYFP was injected into the CN, and AAVretro-pmSyn1-Cre-EBFP was injected into the VAL. This combination allowed selective conditional expression of ChR2-EYFP in neurons that project to the cervical spinal cord but not the VAL. To cover the extent of the VAL, segments C4-T1, and the CN, viruses were injected at locations and depths described above.

### Dual retrograde targeting of VAL and cSC projecting CN neurons to trace axonal projections

To label both VAL projecting and cSC projecting CN neurons, AAV5-EF1a-fDIO-EYFP and AAV1-CAG-FLEX-tdtomato-WPRE were mixed in a 1:1 ratio by volume and injected into the CN. AAVretro-Ef1a-FlpO or AAVretro-pmSyn1-Cre-EBFP were injected into the VAL or cervical spinal cord to label both pathways retrogradely. To cover the extent of the VAL, segments C4-T1, and the CN, viruses were injected at locations and depths described above. To prevent any bias in labeling due to differences in viral efficiency, AAVretro-Ef1a-FlpO and AAVretro-pmSyn1-Cre-EBFP injections were randomized to the VAL or cervical spinal cord in each experimental animal.

### Headplate implantation

In a flat-skull position on a stereotaxic frame, custom-designed Grade 23 titanium alloy headplates (Ti-6Al-4V ELI, South Shore Manufacturing Inc) were placed over the skull, anterior to bregma, using a headplate holder. The headplate was cemented in place using UV-cured dental cement (Tetric EvoFlow, Ivoclar, #595949).

### Optic fiber implantation

For optogenetic stimulation of CN neurons, mice were place in a flat-skull position on a stereotaxic frame, and a fiber optic cannula with a 200 μm core (Doric) was lowered at AP: -1.20 from lambda, ML (right): 1.44, and DV: -2.2 using a stereotaxic manipulator. With the manipulator in place, dental cement was used to secure the fiber to the surface of the skull. The manipulator was then carefully lifted upwards, leaving the fiber in place.

### EMG electrode implantation

EMG surgeries were conducted on animals that had been previously injected with virus and implanted with a headplate and an optic fiber. To implant EMG electrodes, mice were anesthetized, the forelimb and neck were shaved and cleaned with betadine, alcohol, and sterile saline, and the animal was secured in the stereotaxic frame. The skin surrounding the dental cement of the previously implanted headplate and fiber optic was hydrated and gently retracted to the side. The existing dental cement was scored lightly in a criss-cross pattern to create a rough surface to adhere to the EMG electrode. A small incision in the skin was made at the base of the skull, which allowed the EMG electrode leads to tunnel through subcutaneously. A second longer incision was made along the forelimb at the level of the shoulder (just above the shoulder joint) and extending to the forearm (just above the wrist), thus exposing the muscles of interest: biceps brachii (biceps), triceps brachii (triceps), anterior deltoid (ant. delt.), posterior deltoid (p. delt.), and trapezius (trap). Using a hemostat tunneled under the skin between the two incisions, the skin was separated from the underlying muscle, creating a path to lead the EMG wire leads from the connector on the skull to the forelimb muscles. EMG leads were held at the needle end using the hemostat and led under the skin from the neck incision to the forearm incision. At the muscle of interest, the electrode was implanted by inserting the attached needle into the muscle belly, parallel to the longitudinal axis of the muscle. The wire was then pulled through the muscle until the bared electrode region was secured in the muscle. The knot on the electrode remained on the outside of the muscle to secure the electrode at the proximal end, and a secondary double knot was made on the distal end to secure the electrode in place. Any extra wire and the guide needle were then cut off. The mouse was placed in a prone position to implant the trapezius, triceps brachii, and posterior deltoid muscles. To implant the anterior deltoid and biceps brachii, the animal’s nose was tilted ∼45 degrees to the left and the arm was supinated to expose the anterior side of the right forelimb. Once the electrodes were secured in the muscle, the connector was fixed to the scored dental cement using additional cement (C&B Metabond Quick Adhesive Cement System). At the neck incision, the wires leading to the muscles were tucked under the skin and sutured to the neck muscle (6/0 absorbable suture, Covetrus, #57462). Skin incisions were then rinsed with sterile saline, sutured, and cleaned with betadine, alcohol, and sterile saline. Animals were given a subcutaneous injection of buprenorphine-ER (1 mg/kg) and Carprofen (5 mg/kg) and were allowed to recover on a heating pad.

### Tetrode implantation

Animals were anesthetized and implanted with a headplate as described above. A small hole was drilled in the skull using a stereotaxic frame drill attachment ipsilateral (on the right side) to the side of drive implantation. A stainless-steel ground screw was secured into this hole, deep enough to contact the surface of the brain and remain secure in the skull. A craniotomy on the right side of the skull was performed directly over the CN (coordinates from lambda: AP: -1.9-2.9, ML: 1-2 mm), and a section of skull 2 mm in width (ML) and 1 mm in length (AP) was removed. The drive ground wire was wrapped securely around the ground screw in the skull and painted with silver paint (Water-Based Silver Conductive Paint, M.E. Taylor Engineering, Inc, #SP608G) to ensure a low resistance ground connection, left 5-10 minutes to dry, and then secured to the skull with a layer of UV-curing dental cement. Next, the tetrode drive was guided to the coordinates for either the IntA, IntP, or the Med nucleus (see above). The tetrode tip was DV zeroed at the dorsal surface of the brain, a small puncture in the dura was made using a 30-gauge needle, and the drive was slowly inserted to a depth of DV: - 2.00 and left to settle in the tissue for 5-10 minutes. Next, the drive body was secured to the skull and the headplate using dental cement, and custom 3D printed pieces were cemented to the drive to protect the electronic components, forming a nearly complete shield around the drive. A shielding cap was secured to the top of the drive and shielding walls with surgical tape (Dynared Surgical Paper Tape, #K845479), and the animal was removed from the stereotaxic frame, given a subcutaneous injection of buprenorphine-ER (1 mg/kg), and left to recover on a heating pad until awake and mobile.

## Immunohistochemistry and imaging

Animals used for histological purposes were perfused with 10-15 ml cold PBS and 20–25 ml cold 4% paraformaldehyde (PFA) in 0.1M phosphate buffer. Brains and spinal cords down to thoracic vertebrae T2-T4 were dissected and postfixed overnight at 4°C in 4% PFA in 0.1M phosphate buffer. Tissues were then transferred to a 30% sucrose solution for 48 hours before being sectioned on a freezing microtome at 40 μm thickness. Native fluorescence was amplified by immunohistochemistry performed on free-floating sections. Tissue sections were washed in PBS 3 times for 10 min each and blocked in PBS + 5% donkey serum + 0.3% Triton X-100 for 60 min at room temperature. Sections were incubated for 2 days at 4°C in primary antibodies diluted in PBS + 5% donkey serum + 0.3% Triton X-100 at the concentrations listed above. Following primary antibody incubation, sections were washed in PBS 3 times for 10 min each and incubated for 4-12 hrs at 4°C with fluorescent conjugated secondary antibodies diluted in PBS + 5% donkey serum + 0.3% Triton X-100 at the concentrations listed above. Post-secondary antibody incubation, tissue sections were washed in PBS 3 times for 10 min each and then incubated in NeuroTrace diluted in PBS + 0.3% Triton X-100 for 60 min. Sections were then washed in PBS 3 times for 10 min, mounted onto glass slides, and coverslipped with Prolong Diamond or Vectashield mounting media (ProLong™ Diamond Antifade Mountant, ThermoFisher #P36965, VECTASHIELD® Antifade Mounting Medium with DAPI, Vector Laboratories #H-1200-10). To detect axonal fibers, a tyramide signal amplification (TSA) protocol was used, modifying the immunohistochemistry protocol as follows (*68, 69*). After secondary antibody staining, sections were washed in PBS 3 times for 10 min, quenched for 30 min in PBS containing 0.6% H_2_O_2_, and incubated in ABC solution containing avidin-biotin complexes (4 µl solution A + 4 µl solution B to 1 ml PBS-T (PBS + 0.3% Triton X-100), Vector Laboratories, PK-6100). ABC solution was made in advance and allowed to stand for a minimum of 15 min before use. Sections were then washed in PBS 3 times for 10 min and then incubated in tyramide-conjugate Alexa Fluor™ 488 diluted in 0.009% H_2_O_2_ for 30 min. Sections were then washed in PBS 3 times for 10 min, stained with NeuroTrace, and mounted as above. Prepared slides were imaged on a confocal microscope (Zeiss LSM 900; 20X magnification, tiled, z-axis image stacks) or a slide scanner (Olympus VS120; 10X magnification). A motorized stage was used to facilitate the creation of montage images. Images were post-processed in ImageJ, Photoshop (Adobe), and Illustrator (Adobe), and quantification was performed as described below.

## Anatomical characterization of CN networks

### Quantification of retrogradely labeled CN neurons from VAL thalamus and cervical spinal cord

Neuroanatomical analysis of retrogradely labeled CN neurons was performed using a custom CellProfiler pipeline (*70*). Sections beginning at the most anterior end of the CN and ending at the most posterior end were imaged on a confocal microscope and z-stack projections were generated. All images were collected under the same illumination settings. A custom CellProfiler pipeline was developed to delineate CN subnuclei, segment retrogradely labeled nuclei, and colocalize GFP and mCherry-labeled nuclei. CN subnuclei were delineated manually using the Franklin and Paxinos, Mouse Brain Atlas (fourth edition) (*38*), which subdivides the CN into ten distinct sub-nuclei. GFP and mCherry nuclei were segmented using the CellProfiler primary object identification module. Colocalized neurons were then identified using the RelateObjects module. Counts for identified nuclei per subnucleus were then used to quantify subnuclei % distribution and % colocalized neurons.

### Delineation of intersectionally targeted CN networks

Neuroanatomical analysis of INTRSECT axon labeling was performed using the QUINT workflow(*40*), and each brain was processed independently. Brain sections were processed starting at the most anterior portion of the thalamus and ending at the most posterior section of the medulla. All images were batch-processed in ImageJ under a standard brightness and contrast setting to control fluorescence across the dataset. Images were saved as PNGs and high-quality TIFs for atlas registration and axon segmentation, respectively. Brightness and contrast of PNG images could be altered to ensure visualization of neuroanatomical structures during atlas registration, as this does not affect the relative brightness of the files used for axon segmentation.

### Registration

Following pre-processing, each PNG image was registered to the Common Coordinate Framework v3.0; Allen Institute for Brain Science using QuickNII software (*71*). Region identification, tissue size, cutting angle, and spacing of each section were integrated at this step. After initial atlas registration, precise overlay of region borders and section edges were modified individually as necessary using Visualign software. Left and right hemispheric masks were developed in ImageJ for each section. Masking was used to: a) obtain accurate hemispheric quantification of axon tracing; b) correct any atlas inconsistencies due to tissue warping or spacing; and c) remove folds or impurities in the tissue that would register as false positives during pixel quantification.

### Axon segmentation

Labeled axons were segmented using ilastik software (*72*). A random quadrant (∼10% by area) from each TIF image was selected using ImageJ to generate a training dataset, which was imported into ilastik for pixel classification. All pixel features were selected at all provided scales. For images containing fluorescent signals, a single GFP-positive axon was labeled to train the classifier. Images containing no signal were used to define background. Additional labeling of axons was utilized when necessary. Once the segmentation training process was complete, TIF images were batch-processed using the trained workflow to produce simple segmentation PNGs. The PNGs were converted to Glasbey PNGs in ImageJ. Using the quantifier function in Nutil software (NITRC), the final atlas registration files were overlayed with the Glasbey simple segmentation files as well as the hemispheric masks. This workflow produced comprehensive pixel area and object counts for all Allen Brain Atlas regions.

### Data Filtering

For each brain, total object pixels (quantitative measure of axons) were calculated by summing all object pixels across both hemispheres and filtering out object pixels in brain regions that either: a) violated the restrictions set in place during image selection; b) were within the cerebellar injection site (medial, interposed, lateral, infracerebellar, and vestibulocerebellar nuclei); c) were identified as fiber tract structures; or d) received broad region classifications, such as thalamus, cerebellum, medulla, that are further classified into sub-regions, thus avoiding duplicate quantification.

### Data Analysis

Each CN network was quantified based on pixel distribution and pixel density. Pixel distribution was defined as the percentage of total object pixels within a given region relative to total pixels across all regions (object pixels / total object pixels) for a given animal. Pixel density was defined as the percentage of area occupied by object pixels within a given region (object pixels/region pixels). Pixel distribution and pixel density were calculated unilaterally to reflect distinctions in laterality of axon projections.

Four representative brains from CN^VAL^ and CN^VAL/cSC^ targeting were selected based on viral efficacy. For each brain, regions for each hemisphere were sorted by pixel distribution, and the top ten regions within each major brainstem taxonomy structure (interbrain, midbrain, and hindbrain, as defined by the Allen Brain Atlas) were selected. Regions that were identified across all four brains within each CN group and consistently reported a pixel density above 0.1% were used to define the final CN^VAL^ and CN^VAL/cSC^ networks. Regions identified in one group but not meeting the above criteria in the other group were defined as differentially present (marked yellow and green in **Fig. 1F,G**) between CN^VAL^ and CN^VAL/cSC^. The resulting anatomical comparisons were analyzed and visualized in RStudio (version 2024.09.0+375).

CN^VAL^ and CN^VAL/cSC^ schematics (**Fig. 1G**) were designed in Adobe Illustrator (2025). Local distributions of targeted neurons within the CN are qualitatively illustrated for each subpopulation. Pixel densities for each CN group are divided into quartiles and used to determine line thickness to each region, with the thickest lines representing the upper quartile for each CN group. Regions unique to a CN group are shown in color.

## Water reaching behavior

### Behavioral setup

The head fixed water reaching assay was adapted from previously described protocols (*47*). The setup consisted of a removable caddy fitted with custom-built head fixation bars and open rectangular acrylic walls to restrain the animal during the task. A Teensy 3.2 microcontroller utilizing custom scripts was used for experimental control. The water reach system consisted of a calibrated gravity water system (waterspout) gated with a solenoid valve (The Lee Company, #LHDA2433215H), which delivered 5 μL droplets of water reward, a response cue delivered via a tone (#Visaton GmbH & Co. KG, SC 5.9 ND - 8 OHM), a perch sensor (DualLickCmp, HHMI) to detect the animal’s hand at trial initiation, and a waterspout contact sensor (DualLickCmp, HHMI). A reach trial was initiated as follows: the animal was required to continuously place the right hand on the perch sensor for a minimum of 100 ms; if the perch contact was detected, a tone (3000Hz/6000Hz) would signal the start of the trial, and a water reward was presented at the waterspout. The response window was set to a minimum of 1.5 s and a maximum of 3 s in which the animal initiated the trial and contacted the waterspout. Behavioral events (tone onset, reward onset, perch offset, and waterspout contact) were recorded as digital signals using an OpenEphys data acquisition and recording system (Open Ephys Acquisition Board, #SKU: OEPS-9030).

### Behavioral training

Mice implanted with headplates were habituated and trained to reach for the water droplet as follows. Mice were water deprived to a maximum of 85% of their ad libitum body weight for the duration of training and recording. Mice were first gently handled in their home cage to allow the animals to become familiar with the experimenter before being introduced into the behavioral setup. Mice were then habituated to the water reach setup, first by allowing them to explore the caddy freely while intermittently being provided water. Mice were then head fixed to the caddy, initially for ∼5 min per day, progressively increasing over a few days to the maximum recording time of 40 min. Mice were intermittently provided with water during acclimation. To coax mice to begin to use their hands for water retrieval, a small droplet of water was placed under the nose, and the whisker pads were gently stimulated with a blunt needle. This process typically initiates a grooming response, resulting in an encounter with the water droplet, helping animals associate hand movement with reward retrieval. In subsequent training sessions, the waterspout was gradually moved away from the nose until the animals performed a right-handed reach to the spout at a distance of ∼25 mm from the start position (perch) to obtain the reward. Next, the mice were trained to learn the response cue, where the tone and water reward were delivered manually using a button press, with the animals learning to reach only in response to the tone. Once the response cue was learned, animals began to self-initiate reach trials using the automated system by placing their hand on the perch sensor. Behavioral recordings began once the animal consistently reached for the water reward within the contingencies of the experimental paradigm.

### Behavior tracking

Animals were recorded during the reach behavior from three camera angles (horizontally from the side, at a 45-degree angle to the right forelimb, and medio-laterally from the bottom view of the forelimb) to enable 3D kinematic tracking. Videos were acquired at 200 frames per second (fps) using infrared cameras (Basler ace UacA720-520um, Tendelux AI4 IR Illuminator x3) and the pylon Viewer multi-camera acquisition system. Cameras were calibrated using calibration videos of a ChArUco board. In parallel, a frame trigger signal was sent from the cameras to the OpenEphys acquisition system to align video data to neural and EMG recordings and reach events. DeepLabCut was used for keypoint tracking of the hands, digits, nose, mouth, and waterspout (*73*). Anipose was used for 3D calibration and triangulation of tracked data (*74*). The wrist and digits (d1-5, knuckles, midpoint of the digits, and digit tips) of the left and right hand, nose, tongue (tip), lower and upper jaw, and waterspout were tracked. X, Y, and Z coordinates for each tracked keypoint were extracted from Anipose and used for subsequent analysis. For all kinematic analysis of reach trajectories, the knuckle on the third digit of the right hand was used. To account for tracking-related variability in key points, raw data was filtered using an 8^th^-order lowpass Butterworth filter at 50 Hz. Digital events such as the tone, solenoid, perch, waterspout contact, and electrophysiology/ EMG data were aligned to reach kinematics using the camera frame trigger. Custom MATLAB (MathWorks) scripts were created to analyze 3D-kinematics.

### Optogenetic perturbations

Data collection began once the animals consistently reached for the water reward under the task parameters. Each behavioral recording session lasted a minimum of 10 min with ∼45-90 reaches recorded during a single session. Optogenetic stimulation was delivered via pulsed laser (473 nm, 7-8 mW at fiber tip (constant illumination), 18 Hz, 5 ms pulse, 55 ms duty cycle, for 1 second; CNI optical) through an optic fiber implanted over the CN. The laser was controlled via a PulsePal (PulsePal v2, Sanworks LLC, 1102) and operated using the PulsePal MATLAB GUI. Through the Teensy microcontroller, the laser pulse was triggered as the animal initiated the outward phase of the reach, as signaled by offset from the perch sensor. Optogenetic trials were randomized with a 20–25% probability of stimulation, resulting in ∼10-12 stimulated trials per recording session. Stimulation was performed on alternate days of recording to collect sham-stimulated trials and to prevent any potential light-induced changes to reach behavior.

### Reach segmentation

Reaches were first identified using known task contingencies, i.e., 3s of kinematic data after the tone event were extracted and used for further reach segmentation. *Reach start* was identified as follows: From the event-selected reach segment, the maximum speed (speed^max^) was identified as an indicator of the presence of a reaching movement, and the window from tone onset to speed^max^ was used for identification of the reach start. Within this segment, velocity minima were then used to identify initiating submovements. The velocity minimum preceding the submovement that achieved 25% of max reach speed was identified as the reach start. *Reach end* was identified by leveraging the observation that mice increase their grip aperture as the reach nears the target. A window was identified from the time of increased grip aperture to arrival of the hand at the fixed spout position. The velocity minimum closest to the spout contact event and within this window was used to define the reach end. *Speed^max^* was calculated as the maximum of the derivative of instantaneous distance from the target. Reaches that did not cross a minimum threshold of 12 mm of distance traveled were considered incomplete and eliminated from analysis.

### Aligning data to light onset

The time of light onset was recorded as a digital event, and kinematic data were aligned to the recorded event for all light-triggered averaging. To identify a simulated light onset time in light off trials, the time of delay of light onset after reach start was calculated in light on trials, and the median within each animal was used to identify a similar timepoint in light off trials for each trial.

### Kinematic quantification

#### Acceleration peaks per mm

To quantify directional changes in movement induced by the light, acceleration peaks were used as a proxy for abrupt changes in movement. Acceleration peaks greater than 1000 mm/s^2^ were quantified between the reach start and reach end and normalized to the Euclidean length of the reach.

#### Latency to first acceleration peak after light onset

To identify the latency of light-induced directional changes in movement, peaks in absolute jerk, which represent an abrupt change in acceleration, were identified after light onset and compared to a similar timepoint in light off trials. The latency in milliseconds to the first peak is reported.

#### Direction of light-triggered movements

A light-triggered movement (LTM) was defined as the movement after the first stimulation pulse until the next break in the movement, defined as a local velocity minimum. To compare to light off trials, submovements were defined as segments of the reach between two velocity minima, and the direction vectors of the first submovement in light off trials were compared to the first LTM in light on trials. To assess the direction of light-triggered movements during inward phases of movement, stimulation pulses that landed at time points where the hand was moving inward for at least 15 ms were identified. Submovements triggered between the light pulse and 55 ms later, when the next light pulse was delivered, were used to quantify movement direction. For comparison to light off trials, timepoints in the light off reach that matched light on data in 3D position, 3D velocity, and percent of reach progress were identified, and submovements in the subsequent 55 ms were used for analysis. The dot product of the submovement unit vectors and the AP/DV axis vector was calculated and quantified using hierarchical bootstrapping.

#### Relative change in velocity

To quantify changes in velocity following optogenetic stimulation, the change in AP and DV velocity relative to velocity at light onset was calculated for each stimulation trial, up to 55 ms after the first stimulation pulse. For light off trials, the change in AP and DV velocity was calculated relative to the sham light onset time calculated as above. Within animal means in a window 20–25 ms post light stimulation or sham were then averaged across animals to generate across-animal plots. 20–25 ms was chosen based on latency analysis of the onset of a change in acceleration described above, which occurs at ∼16–19 ms (see **Fig. 2**).

#### AP displacement

To quantify the relative displacement in the forward vs backward direction per trial, the ratio of forward and backward displacement was calculated. Forward displacement was defined as the area under the curve of the positive component of the velocity profile in the AP dimension, and backward displacement was defined as the area under the curve of the negative component of the AP velocity profile.

## Electrophysiological recording of CN neurons

### Recording

Animals with chronically implanted tetrode drives were secured in the head fixation caddy, the shielding cap was removed, and an Intan 32-channel RHD head stage (RHD 32-Channel Recording Headstages, Intan, #C3314) was plugged into the drive via an Omnetics connector. The head stage was connected to an OpenEphys acquisition board via an SPI cable (RHD 3-ft (0.9 m) standard SPI cable, #C3203, RHD 1-ft (0.3 m) ultra-thin SPI cable, #C3211). The caddy was then inserted into the recording rig, and perch contact sensor cables were plugged in. Neural data were recorded at 30 kHz and displayed in the Open Ephys GUI. Filtered extracellular voltage traces across all channels were visualized to determine if any units were present on any tetrode bundles. If at least one unit was visible on multiple channels within a bundle, the recording session proceeded. If no units were visible, the session was ended. At the end of each session, the head stage was unplugged, and the drive screw, which controls the vertical position of the fiber in the brain relative to the skull, was turned clockwise ¼ turn, corresponding to 32.5 μm of ventral travel. The shielding cap was reattached to the drive shielding wall via surgical tape, and the mouse was removed from head fixation and returned to its home cage. Recording sessions were collected starting from the first session where units were visible on the Open Ephys GUI until the drive was estimated to be ∼0.25 mm ventral to the implanted CN, according to calculated drive position (current drive position = implanted depth + drive screw rotations X vertical travel of single rotation). After the termination of recording sessions, animals were perfused as described above, and before the brain was explanted, small electrolytic lesions were created at the tip of each tetrode bundle by passing 100 mA of current for <5 s between two wires of each bundle. Brain tissue was then sectioned, mounted, and imaged on an Olympus VS120 slide scanner. Putative final tetrode locations and depths were determined by matching sections to mouse coronal atlas images using image analysis tools in FIJI (*38, 75*). From this final location, each preceding recording session location was calculated by subtracting from the probe depth each session’s vertical travel distance. Only sessions in which all the bundles were contained within CN were used for further analysis.

### Data analysis

Raw electrophysiology data were processed in the Python-based SpikeInterface platform (*76*). First, voltage signals were filtered with a 300 Hz low pass and 5000 Hz high pass followed by common median reference filters. Filtered data were then spike sorted using Kilosort2 (https://github.com/MouseLand/Kilosort), and units were manually curated in Phy2 (https://github.com/cortex-lab/phy). Only well isolated units fulfilling quality metrics (refractory period contamination ratio < 0.1, mean firing rate > 1 Hz) and having a physiological waveform shape that was present throughout the whole session passed curation and were imported into MATLAB for further analysis.

Import of spike clusters and quality metric data from Spike Interface and Phy into MATLAB was assisted by the “spikes” repository (https://github.com/cortex-lab/spikes). Import and filtering of kinematic data from Anipose and event data from Open Ephys was performed using in-house MATLAB scripts(*74*). To determine the average activity of recorded units during reaching, for each unit, histograms of spike counts across all reaches centered on peak reach speed with a bin width of 1 ms were generated, normalized by reach count, and smoothed with a 10 ms Gaussian. This reach-averaged firing rate was then z-scored using the mean firing rate and standard deviation of firing rate of that unit. These quantities were calculated from 1000 randomly sampled windows from the session. Instantaneous firing rates were calculated by taking the inverse of the inter-spike intervals, convolving with a 10 ms kernel Gaussian, and then resampling at 200 Hz to match the sampling rate of the kinematic data.

### Cross-correlation analysis

To determine the relationship between unit activity and hand forward velocity on a per unit basis during reaching, instantaneous firing rates of units were cross-correlated with the AP component of velocity. A window from 100 ms before max speed to 150 ms after max speed was selected, as this time range contained reach initiation, the major acceleration and deceleration phases, and movement completion for most reaches.

For each reach, Pearson correlations were computed between velocity at frame *t* and firing rate at frame *t + L* for lags *L* spanning -20 to +20 frames (±100 ms at 200 fps); under this convention negative lags indicate firing rate preceding velocity. Correlations were Fisher *z*-transformed, aggregated across reaches by the median, and back-transformed, 1,000 bootstrap resamples of reaches drawn with replacement. Because firing rate and velocity both follow a stereotyped reach profile, a trial-shuffled control was computed identically within each resample with firing-rate traces re-indexed to break their correspondence with velocity; the test statistic was the shuffle-subtracted correlation (real minus shuffled), averaged across bootstraps.

Significance was assessed by cluster-based permutation (*77*), with family-wise error controlled across the lag axis by the maximum-statistic method (*78*). A null distribution was generated from 10,000 permutations in which reach labels of the firing-rate matrix were permuted globally. Positive and negative clusters were detected separately. For each direction, a cluster-forming threshold was set per lag at the 95th percentile of the null at that lag, contiguous above-threshold runs were retained as candidate clusters, and cluster mass was defined as the summed statistic across the cluster. Within each permutation, the largest mass across both directions was retained, yielding a null distribution of maximum cluster mass per unit. Each observed cluster received a corrected p-value equal to the proportion of permutations whose maximum mass equalled or exceeded it, with significance defined as p < 0.05. Exceeding the cluster-forming threshold determined cluster membership only and did not constitute a test at individual lags.

Clusters within a unit were non-overlapping in lag by construction. Cluster center lag was defined as the mass-weighted center (centroid) of lags within the cluster. Units with a significant positive cluster centered at a lag less than 0 ms were labelled ‘AP+’, units with a significant negative cluster centered less than 0 ms were labelled ‘AP-’, and units with no significant cluster at any lag were labelled ‘AP uncorrelated’.

### Sliding window correlation analysis

To measure the relationship between firing rate and AP velocity continuously across the reach, we computed a between-reach correlation at each time point relative to maximum speed. For a unit with cluster lag *L* (where negative indicates firing rate precedes velocity change), at each time *t* from -100 ms to +150 ms relative to maximum speed, firing rate at *t + L* was correlated across reaches with AP velocity at *t*, yielding a correlation time course per unit. Units without significant correlations (AP uncorrelated) were assigned a lag drawn with replacement from the AP+/AP-lag distribution. Traces were z-scored and smoothed for visualization only. Significance was assessed per unit by permutation. The pairing between firing rates and velocities was shuffled 10,000 times, recomputing the correlation time course each time, with the threshold set at the 95^th^ percentile of the maximum correlation time courses across shuffles, effectively controlling family-wise error across time with the maximum statistic (*78*).

### Mutual information analysis

To quantify the dependencies (both linear and nonlinear) between AP velocity and unit firing rate of AP+, AP-, and AP uncorrelated units using an information theoretic approach, we applied a Kraskov, Stögbauer, Grassberger (KSG) mutual information estimator (*48*). The same data used for cross-correlation analyses above were used as input to the estimator, with estimations repeated at lags from -100 to 0 ms, generating a lag-mutual information curve for each unit. The peak mutual information and corresponding lag were extracted for comparison to max/min cross-correlation and lag. This process was repeated with the trial-shuffled conditions for each of the three groups of units, and peak mutual information for each group was compared to its trial-shuffled control with a bootstrap comparison test and Holm-Bonferroni correction for multiple comparisons (see hierarchical bootstrap analysis below).

### Hierarchical bootstrap analysis

Units were nested within sessions within mice, so unit-level statistics were not independent. We therefore estimated confidence intervals and p-values using a three-level hierarchical bootstrap (*79*). On each iteration, mice were resampled with replacement, then sessions within each resampled mouse, then units within each resampled session, each level resampled to its observed size. The statistic of interest was recomputed on each resampled set, and 10,000 iterations were run. Confidence intervals are 95% percentile intervals of the resulting distribution; p-values are two-sided percentile p-values, defined as twice the smaller tail proportion relative to the null value and floored at 1/n iterations, so a reported p < 10⁻⁴ indicates that no iteration crossed the null. Where a family of related tests was conducted, p-values were Holm-corrected within that family. This procedure was applied to the mutual information estimates and their trial-shuffled controls.

### Firing rate quartile analysis

To visualize the relationship between firing rate and subsequent kinematics, unit spike times and AP velocity were aligned to a given event (reach start, maximum speed, a late reach point when velocity fell below 130 mm/s, or reach end). Reaches were divided into quartiles by mean firing rate over a 50 ms window relative to the event, and mean firing rate and AP velocity were plotted per quartile with SEM as shaded region. Quartiles are for visualization only; the analysis below uses individual reaches.

### Firing rate-velocity regression

For each unit, the mean firing rate over the quartile window per reach was paired with AP velocity sampled at that unit’s cross correlation cluster lag relative to the quartile window center (for example, firing rate taken at center of quartile window and AP velocity taken 55 ms later for a unit with a correlation lag of -55 ms). An ordinary least-squares line was then fitted across reaches.

## EMG recordings

### Recording

Animals were placed in the water reach setup, and the electrodes were connected to the acquisition system via a custom-built EMG connector. Bipolar EMG signals were recorded at 30 kHz, amplified first via differential preamplifiers (ultra-low noise preamplifier, University of Cologne, Model: MA 103), bandpass filtered at 100-5000 Hz, amplified again (University of Cologne, Model: MA 102), digitized, and acquired using the OpenEphys acquisition system. Impedance measurements were taken during every recording session, and electrodes were considered intact if impedance measurements were in the range of ∼10-100 kΩ. Electrode impedance was found to be stable throughout the duration of a recording session. In the case of electrode damage, impedance measurements increased to 1-3 MΩ, and these recordings were not used for subsequent data analysis.

### Data Analysis

Raw EMG data was first filtered using a 4^th^-order bandpass Butterworth filter with a frequency cutoff between 200-1000 Hz. EMG signals were then rectified and filtered using a 4^th^-order lowpass Butterworth filter with a frequency cutoff of 50 Hz. EMG data was aligned to tone and reach events (start, end, and speed^max^) for subsequent analysis. Custom MATLAB scripts were created to analyze muscle activity relative to light onset and kinematic data.

### Analysis of kinematic and EMG similarity

For unstimulated reaches, three-dimensional forelimb trajectories and corresponding EMG profiles were time-normalized and matched by trial identity. Kinematic similarity was quantified pairwise for every trial using root mean square error (RMSE) between their 3D trajectories, while EMG similarity for the same pair of reaches was quantified using cosine distance between the concatenated muscle-activity profiles. Pairs below the 10th percentile of RMSE were classified as highly kinematically similar, whereas pairs above the 90th percentile were classified as kinematically dissimilar. EMG cosine distance distributions were summarized within each animal using their median and measures of variability, including range, IQR, and coefficient of variation. For light-triggered movements, kinematic similarity and EMG similarity were calculated as above in a 40 ms time window after light onset, and median, range, IQR, and coefficient of variation were measured in the most kinematically similar and dissimilar trial pairs.

### Latency of light-triggered muscle activity

To identify muscles with a light-induced increase in muscle activity, peaks in z-scored muscle activity one standard deviation above the mean for each muscle were calculated. Two criteria were used for identifying muscles with short-latency effects: a) a peak probability >70% within 20 ms of light onset; and b) standard deviation of the latency of identified peaks < 4 ms.

## Electrophysiological recording of spinal motor neurons

Preparations for the experiments, including anesthesia, tracheotomy, pneumothorax, artificial respiration, laminectomy, and craniotomy, were performed following previously published procedures (*37, 80*).

### Stimulation and recording

The ipsilateral deep radial nerve (DR) was stimulated with needle electrodes inserted into the forelimb for guidance to identify forelimb-innervating motor neurons and the ventral horn antidromically and to assess the physiological integrity of the spinal cord. The CN were stimulated with a tungsten electrode (100 KΩ impedance, 25° forward angle, 20-100 μA) inserted into the Int at 2.7-3.5 mm rostral to obex, 1.4-2.0 mm ipsilateral from the midline, and 1.5 mm ventral from the surface of the cerebellum.

Intracellular motor neuron recordings were conducted by using sharp borosilicate glass micropipettes (10–20 MΩ impedance) filled with 2 M potassium citrate, pH 7.4. To measure the effect of CN stimulation on forelimb-innervating motor neurons (N=5 mice), the glass micropipette was inserted into C6/C7 at 10-12° lateral from the vertical line to a depth ranging from 0.8 to 1.2 mm from the surface of the cord toward laminae IX. The DR motor neurons were identified by antidromic activation after DR stimulation at 5xT. We also included unidentified motor neurons recorded in the same tracks as DR motor neuron recordings. Cord dorsum potentials were recorded with a silver ball electrode on the surface of the spinal cord near the recording site to monitor the incoming volley and to assess the physiological integrity of the spinal cord.

### Statistics and data collection

Mice from each litter were randomly allocated to different groups for the electrophysiological and behavioral experiments. Group sizes were not predetermined, but sample sizes were comparable to those commonly used in similar experiments and were selected to allow for the use of appropriate statistical tests. For water reach kinematics, data collection and analysis were automated, minimizing any potential influence by the experimenter. Animals were only excluded from behavioral experiments if data collection could not be performed, viral expression was not detected in the CN after amplification of fluorescent signals, or viral expression was determined to be inefficient in intersectional targeting of each population. These cases were: two animals in the CN^VAL^ group and one animal in the CN^VAL/cSC^ group that did not show expression of virus; one animal in the CN^cSC^ group that showed above 0.1% axonal pixel density in the VAL thalamus, indicating inefficient intersectional targeting; one animal in the CN^cSC^ group that did not reliably perform the behavior during training; two animals in the CN^cSC^ group and one animal in the CN^VAL^ group that pulled out their optic fiber or EMG implants; and one animal in the CN^VAL^ group whose recording data could not be aligned to video data due to experimental error. Results are shown as individual values with median, 25th and 75th percentiles indicated, or box- and-whisker plots indicating the median, 25th and 75th percentiles, and range, unless otherwise indicated. Normality was assessed using the Shapiro-Wilk test. All statistical tests and N values for each experiment are indicated in the figure legends. For kinematic data, statistical analysis was performed in one of two ways: within-animal paired t-tests of summary statistics between conditions, and joint probability analysis for bootstrapped distributions. For paired t tests, p < 0.05 was considered significant, * indicates p < 0.05, ** < 0.01, *** <0.001, **** < 0.0001. To calculate significant differences between two bootstrapped distributions, the joint probability distribution of the bootstrapped medians between the two groups was computed (*79, 81*). Probability < 0.05, was considered significant. All p-values are indicated in the figure legends. Statistical analysis was performed in MATLAB or Prism (version 10.2.3, GraphPad).

## Acknowledgements

We are grateful to Phong Nguyen for assistance with mouse husbandry, histology, and lab operations; Aakash Bhegade, Madelyn Peck, Zoe Sarafis, Garrett Tan, James Taniguchi, and Elaine Zhao for help training models for limb tracking; Steve Barry at the Salk Institute Machine Shop for help with device fabrication; Ed Callaway for the pAAV-hSyn-H2B-GFP and pAAV-hSyn-H2B-mCherry plasmids; and Amy Cao for assistance with the summary figure. We thank Tejapratap Bollu, Denis Jabaudon, Boaz Styr, John Tuthill, Michael Yartsev, and members of the Azim lab for valuable discussion and comments on the manuscript.

## Funding

National Institutes of Health grant CCSG P30 CA014195 (GT3 Core Facility of the Salk Institute; RRID:SCR_014847)

National Institutes of Health grant CCSG P30 CA014195 (The Waitt Advanced Biophotonics Core Facility of the Salk Institute; RRID:SCR_014838)

San Diego Nathan Shock Center National Institutes of Health grant P30 AG068635 (The Waitt Advanced Biophotonics Core Facility of the Salk Institute; RRID:SCR_014838)

The Henry L. Guenther Foundation (The Waitt Advanced Biophotonics Core Facility of the Salk Institute; RRID:SCR_014838)

The Waitt Foundation (The Waitt Advanced Biophotonics Core Facility of the Salk Institute; RRID:SCR_014838)

Salk Alumni Fellowship Award (AT)

Salk Women & Science Research Award (AT)

National Institutes of Health grant F31 NS130972 (OW)

Swedish Research Council grant 2015-03581 (JJ)

National Institutes of Health grant F32 NS126231(KWH)

Salk Alumni Fellowship Award (KWH)

Human Frontier Science Program Postdoctoral Fellowship (MET)

National Institutes of Health grant F32 NS120998 (AN)

Salk Pioneer Fund Postdoctoral Scholar Award (AN)

National Institutes of Health grant R01 DK134857 (AIC)

National Institutes of Health grant R01 DK124801 (AIC)

National Institutes of Health grant R01 NS124844 (AIC)

National Institutes of Health grant RF1 NS128898 (EA, AIC)

National Institutes of Health grant U19 NS112959 (EA)

National Institutes of Health grant DP2 NS105555 (EA)

National Institutes of Health grant R01 NS111479 (EA)

Searle Scholars Program (EA)

The Pew Charitable Trusts (EA) The McKnight Foundation (EA)

The Brain Research Foundation (EA) The Callahan Foundation (EA)

## Author contributions

Conceptualization: AT, AIC, EA

Experimental design: AT, OW, EA

Viral injections: AT

Anatomical experiments, behavioral experiments, and EMG recordings: AT, ER of kinematic and EMG data: AT

Analysis of anatomical data: ER

Development of anatomical analysis pipelines: OW

CN electrophysiological recordings and data analysis: OW, AT

Motor neuron electrophysiological recordings and data analysis: JJ

Tetrode microdrive design and development of head fixed water reaching task, electrophysiology recording system, and water reaching data processing pipelines: KWH

Assistance with spike sorting: EB

Assistance with anatomical and behavioral experiments: ER, RY, EB

Fabrication of EMG electrodes and assistance with anatomical data analysis: DS

Assistance with surgery and establishing electrophysiological recording setup: MET

Preliminary anatomical data: JMC

Consultation on EMG data analysis: AN

Funding acquisition: EA, AIC

Project administration and supervision: EA

Materials and Methods text, results text, and figures: AT, OW, ER, EA

Writing – original draft: AT, EA

Writing – review and editing: AT, OW, ER, JJ, KWH, AN, JMC, AIC, EA

## Data and materials availability

All data are available in the manuscript or the supplementary materials.

## List of Supplementary Materials

Supplementary Text Figs. S1 to S12

Supplementary Tables S1 to S2

**Movie 1. Normal movement execution during the head fixed water reach task.** Example animal reaching toward a waterspout without optogenetic stimulation. Videos are captured from the side (top), underneath (middle), and angled at 45 degrees (bottom). Automated 2D markerless tracking of the right hand and 3D reconstruction were used for kinematic quantification (fps, frames per second).

**Movie 2. Activation of CN^VAL^ neurons drives inward limb movement.** CN^VAL^ neurons were targeted using AAV-mediated intersectional expression of ChR2-EYFP. Photostimulation (473 nm, 18 Hz, 5 ms pulse width) resulted in submovements shortly after each light pulse primarily directed toward the body (most visible in the top view). See **Fig. 2**.

**Movie 3. Activation of CN^VAL/cSC^ neurons drives outward limb movement.** CN^VAL/cSC^ neurons were targeted using AAV-mediated intersectional expression of ChR2-EYFP. Photostimulation (473 nm, 18 Hz, 5 ms pulse width) resulted in submovements shortly after each light pulse primarily directed away from the body (most visible in the top view). See **Fig. 2**.

**Movie 4. Activation of CN^cSC^ neurons does not drive directional limb movement.** CN^cSC^ neurons were targeted using AAV-mediated intersectional expression of ChR2-EYFP. Photostimulation (473 nm, 18 Hz, 5 ms pulse width) did not elicit submovements (most visible in the top view). See **fig. S4**.

## Supplementary Text

We attempted to link the anatomically defined CN^VAL^ and CN^VAL/cSC^ groups to the electrophysiologically defined AP+ and AP-classes through two sets of optotagging experiments. First, using the optetrode recording approach (**Fig. 3, fig. S9**), we performed broad labelling of thalamus-projecting CN neurons (retrograde AAV-Cre injected into VAL thalamus of transgenic animals conditionally expressing the excitatory opsin ChR2 or the inhibitory opsin GtACR1). We pursued this approach, which required the use of only one virus, to assess the feasibility of optotagging from a large population of CN neurons. Importantly, this strategy does not allow us to distinguish CN^VAL^ and CN^VAL/cSC^ neurons. We recorded from 11 mice each performing 3-10 behavioral sessions, adjusting the optetrode location on the z-axis using a microdrive each session to target new units. We were only able to identify a handful of units that were putatively excited by ChR2 activation from this broad population. Moreover, GtACR1 tagging introduced difficult confounds due to the tendency of GtACR1+ neurons to spike after light onset, the difficulty of establishing precise onset and jitter of the silencing effect, and variable effects on spiking throughout the course of photostimulation. With very few units and the inability to distinguish CN^VAL^ and CN^VAL/cSC^ neurons in this experiment, we decided against continuing to pursue this recording approach.

In a separate set of experiments, we performed acute recordings of CN neurons using silicon probes with attached tapered lambda optical fibers, in an attempt to increase yield. Using our triple-viral intersectional strategy, we expressed ChRmine (a red-shifted opsin that allows increased light penetration) in CN^VAL^ and CN^VAL/cSC^ neurons. We recorded from 10 animals each performing 3-5 sessions, with the probe implanted in a slightly different location within Int and Med nuclei every session. In total, we were able to identify 232 well-isolated CN units. Using cross-correlation analysis (as in **Fig. 3** and **fig. S9**), we found 25 units with significant positive and 32 units with significant negative correlations with AP velocity (across negative and positive lags). From these units, we found several putatively light-responsive neurons (13 CN^VAL^ neurons and 5 CN^VAL/cSC^ neurons across all animals) using standard tagging criteria (assessed with the Stimulus-Associated Latency Test (SALT), latency < 10 ms after light onset, coefficient of variation < 0.55, reliability > 0.5). However, close inspection of rasters aligned to light onset and spiking jitter (measured as SD of latency) revealed complex light induced responses that suggested most might not be *bona fide* tagged neurons but rather could be synaptically activated units. For example, many units had relatively long response latencies and jitter values were quite large, ranging between 3 – 5.5 ms, far above a typical jitter of < 2 ms that would be expected from a tagged neuron. We therefore used more stringent criteria that better align with the large single-channel conductance and ultra-high photosensitivity of ChRmine (latency < 5 ms and jitter < 2 ms). Applying these criteria, we found only one convincingly tagged CN^VAL^ unit. Notably, this unit falls squarely within the AP-category with a negative lag, as predicted for a class of neurons that promotes inward limb movement. While these results provide initial evidence that CN^VAL^ neurons might belong to the AP-class, we would require at least an order of magnitude more mice to tag enough CN^VAL^ neurons to make a statistically valid claim. Moreover, given their greater sparsity, an even larger number of mice would likely be needed to tag CN^VAL/cSC^ neurons.

The primary challenge here is one of yield. Targeting CN^VAL^ and CN^VAL/cSC^ neurons requires the injection of three different AAVs into three distinct locations in the nervous system. The total efficiency of targeting is progressively limited by the convergent expression of all three viruses. Our findings highlight the need for more efficient approaches to link anatomically defined CN classes to their electrophysiological activity during limb movement.

**fig. S1.**
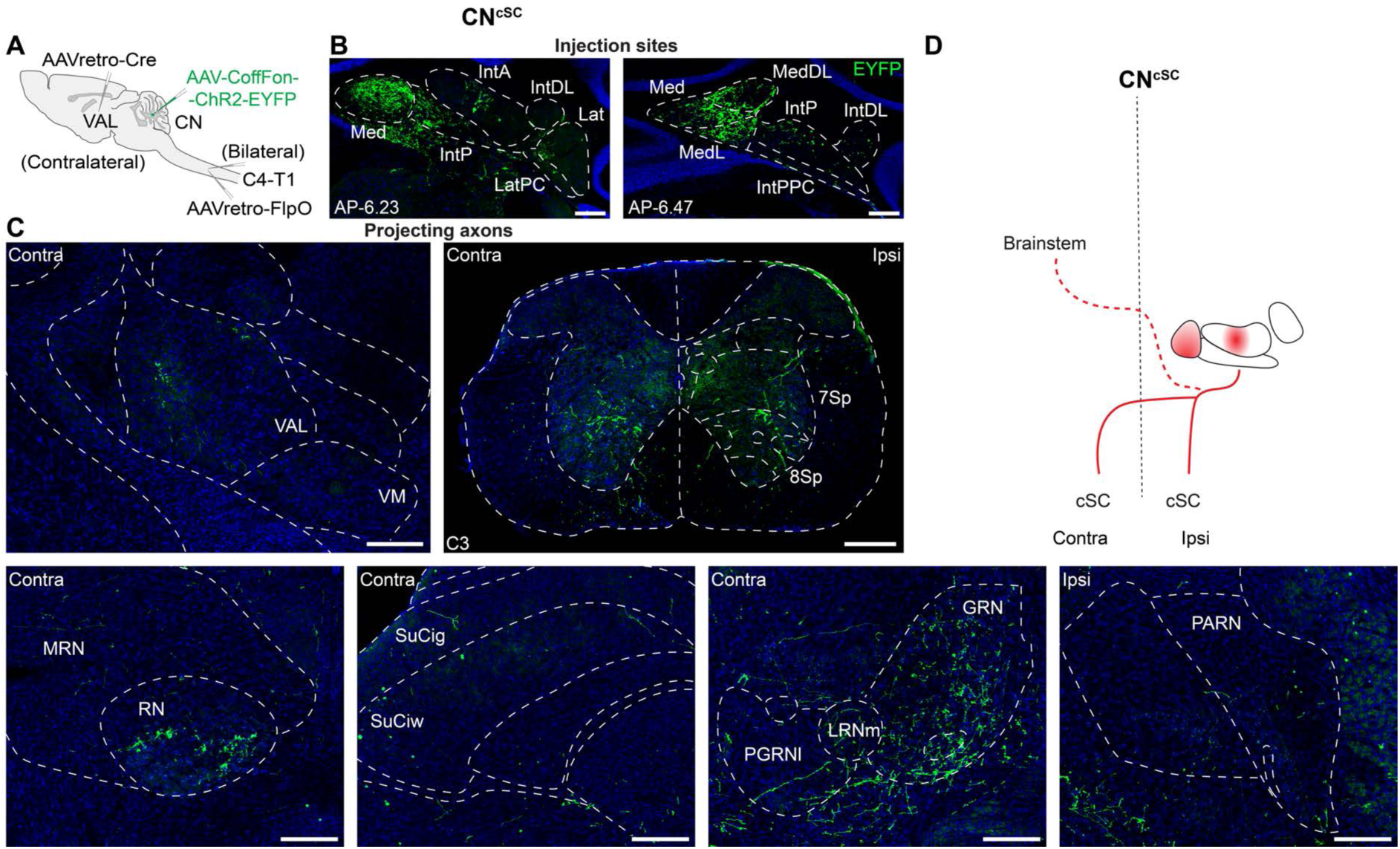
Intersectional targeting of CN^cSC^ neurons. **(A)** Strategy for targeting CN^cSC^ neurons. **(B)** Expression of ChR2-EYFP in the CN at the injection sites. AP coordinates indicate distance from bregma. **(C)** ChR2-EYFP expression in axons with minimal projections to the VAL (top left) and more extensive projections to the cSC (top right). Some relatively sparse projections (when compared to the other CN subpopulations) can also be seen in other brain targets; however, these regions do not cross thresholds applied for CN^VAL^ and CN^VAL/cSC^ neurons (Materials and Methods). Scale bars: 250 μm. Brain region abbreviations are defined in **table S1**. **(D)** Schematic of CN^cSC^ outputs showing bilateral targeting of the cervical spinal cord and sparse projections to the brainstem.

**fig. S2.**
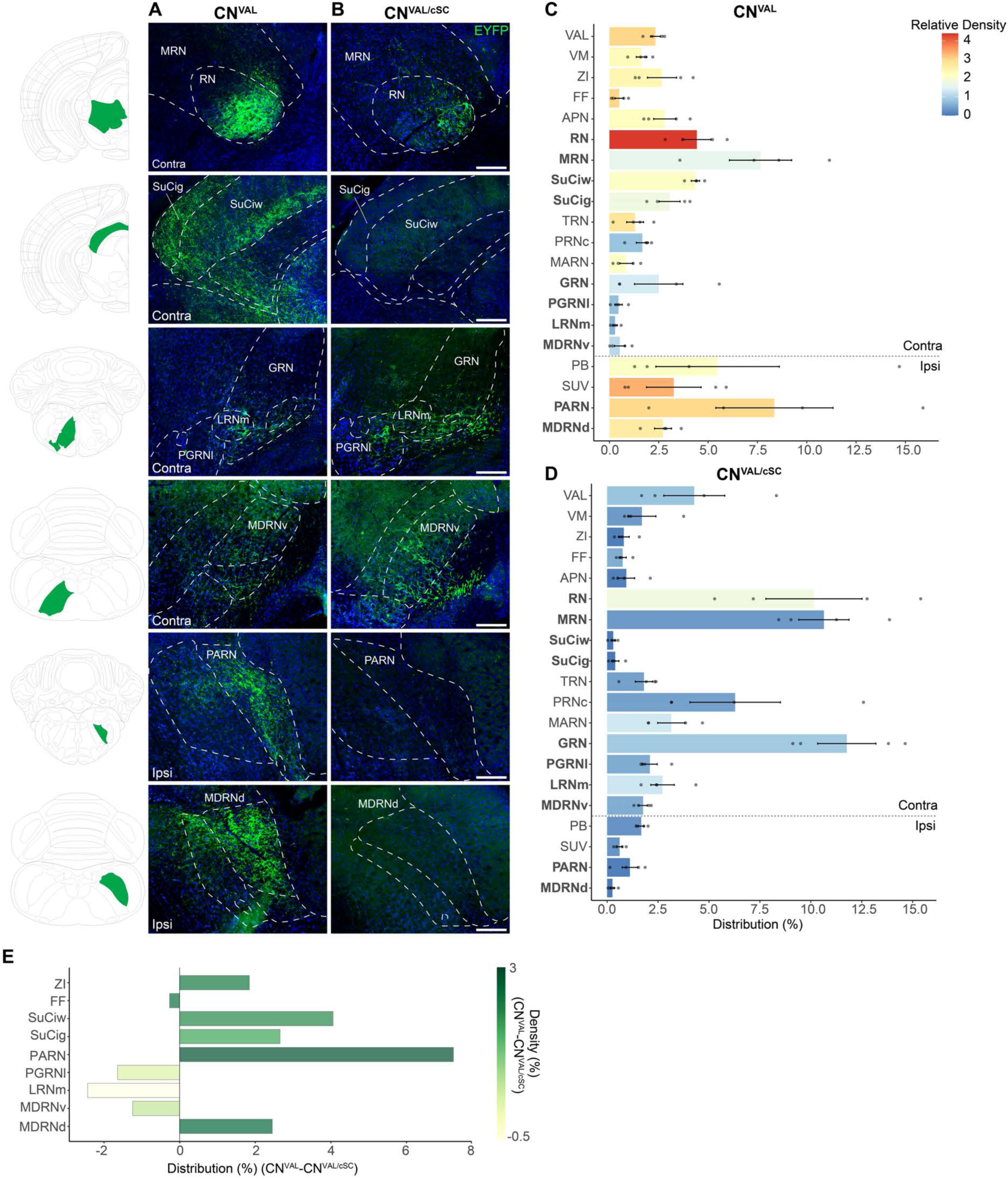
Subcortical targets of CN^VAL^ and CN^VAL/cSC^ neurons. (A,B) Axonal projections of CN^VAL^ (A) and CN^VAL/cSC^ (B) neurons to brainstem targets. Regions of interest are shown in order of their rostral (top) to caudal (bottom) position. Representative atlas images (left) show regions of interest in green, highlighting differences in projection targets (see **Fig. 1F,G**). Scale bars: 250 μm. **(C,D)** Distribution of axonal projections across brain regions expressed as a percentage of total pixels within each animal (mean ± SEM), and density (based on pixels/region area; indicated by the heatmap) of labeled axons in CN^VAL^ (C) and CN^VAL/cSC^ (D) animals across the brain (CN^VAL^, N=4; CN^VAL/cSC^, N=4). Bold labels are regions depicted in (A,B). **(E)** Delta between axon terminal distribution and density in projection targets that are differentially labelled between CN^VAL^ and CN^VAL/cSC^ (see Fig. 1F,**G**). Brain region abbreviations are defined in **table S1**.

**fig. S3.**
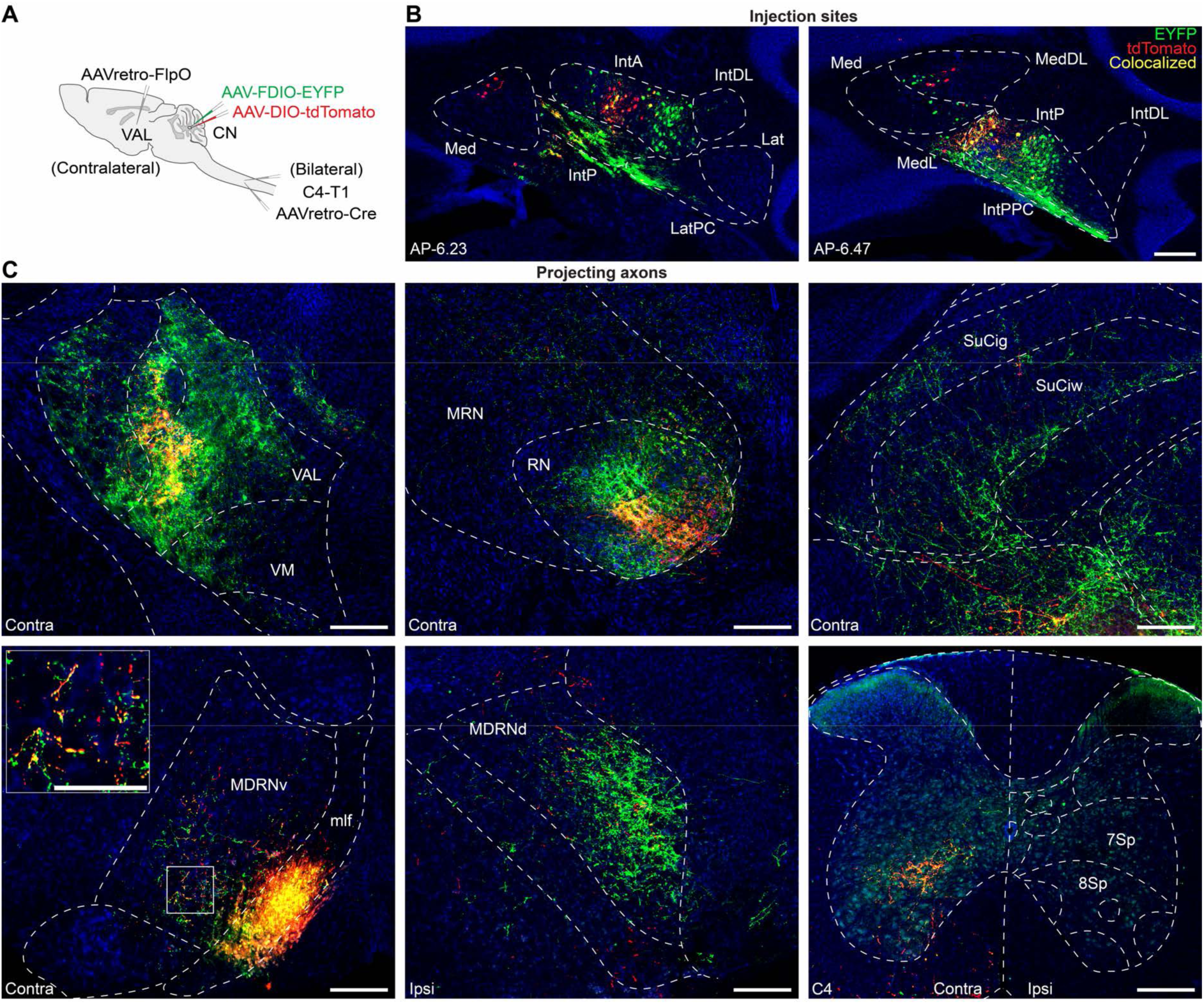
Spatial organization of putative CN^VAL^ and CN^VAL/cSC^ axon projections in downstream targets. **(A)** Strategy for dual labeling of VAL and cSC projecting neurons and their axonal projections. **(B)** Expression of EYFP and tdTomato in the CN at the injection sites. Putative CN^VAL^ (EYFP+, green), CN^VAL/cSC^ (EYFP+/tdTomato+, yellow), and CN^cSC^ (tdTomato+, red) neurons can be seen. **(C)** Axon projections of labeled neurons in the VAL and RN show that CN^VAL/cSC^ axons occupy a subregion of the overall CN^VAL^ projection field (top left and middle). Projections to the SuC (top right) and MDRNd (bottom middle) are predominantly from CN^VAL^ neurons (green), representing regions where the pathways diverge. Projections to the MDRNv (bottom left; inset shows higher magnification) and cSC (C4; bottom right) show colabeled axons (yellow) spread across the overall CN projection field, indicating that CN^VAL/cSC^ neurons contribute significantly to projections to these areas. Scale bars: 250 μm; inset, 125 μm. N=3. Brain region abbreviations are defined in **table S1**.

**fig. S4.**
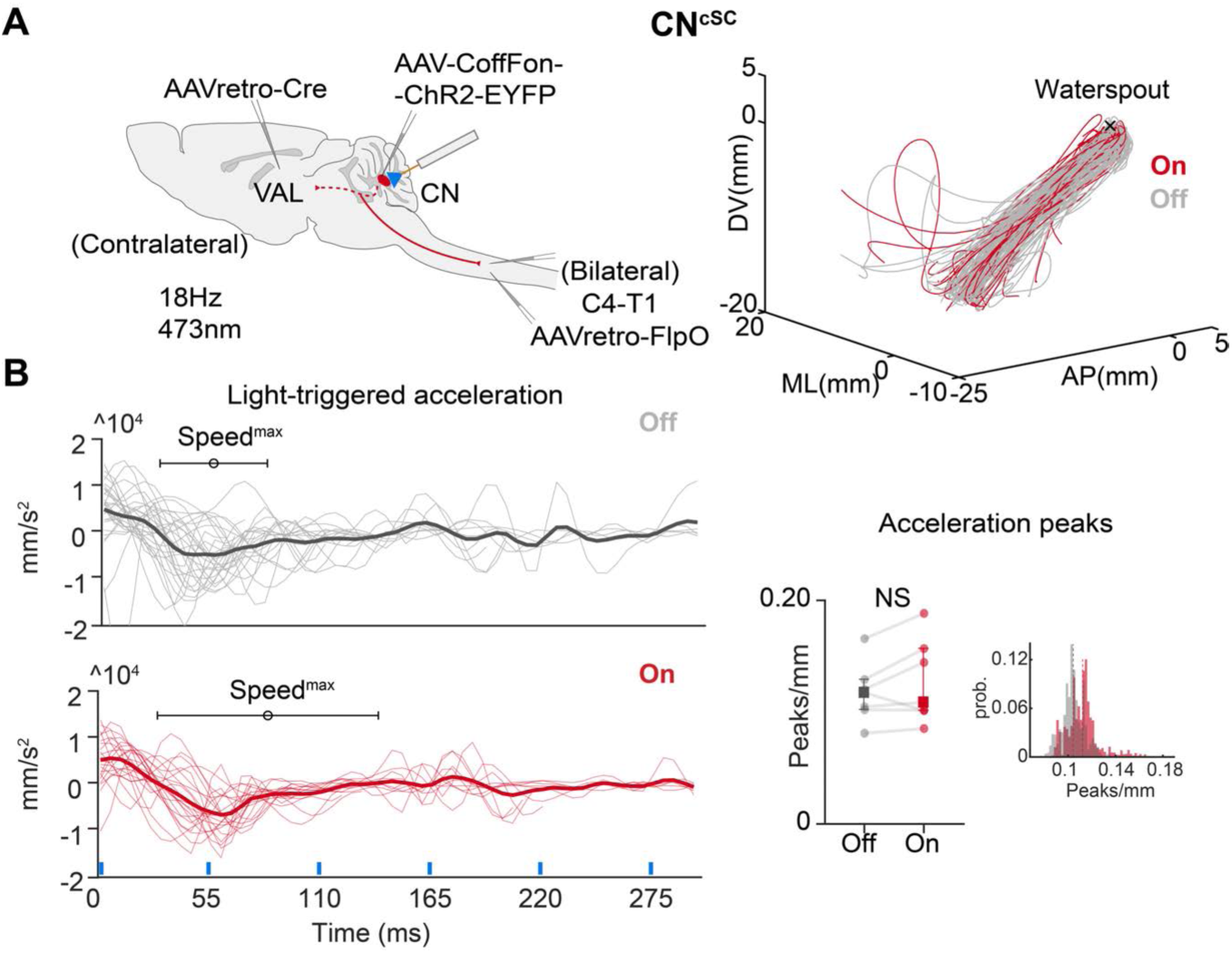
Activation of CN^cSC^ neurons does not evoke directional limb movements. **(A)** Intersectional strategy to target CN^cSC^ neurons for expression of ChR2-EYFP (left). 3D trajectories of all reach trials from an example animal; grey lines indicate light off trials, red lines indicate light on trials (right). **(B)** Hand acceleration triggered at light onset in CN^cSC^ trials (red) compared to similar timepoints during light off trials (grey) from a representative animal; thicker lines represent mean, thinner lines represent single trials, blue lines indicate pulses of light, distributions of when Speed^max^ occurred are shown (mean ± SEM) (left). Number of acceleration peaks normalized to path length across animals (right; N=7; NS, p=0.1687, paired t-test); circles represent averages within animal, squares and error bars represent median and IQR across mice. Hierarchical bootstrapped distribution of medians (inset).

**fig. S5.**
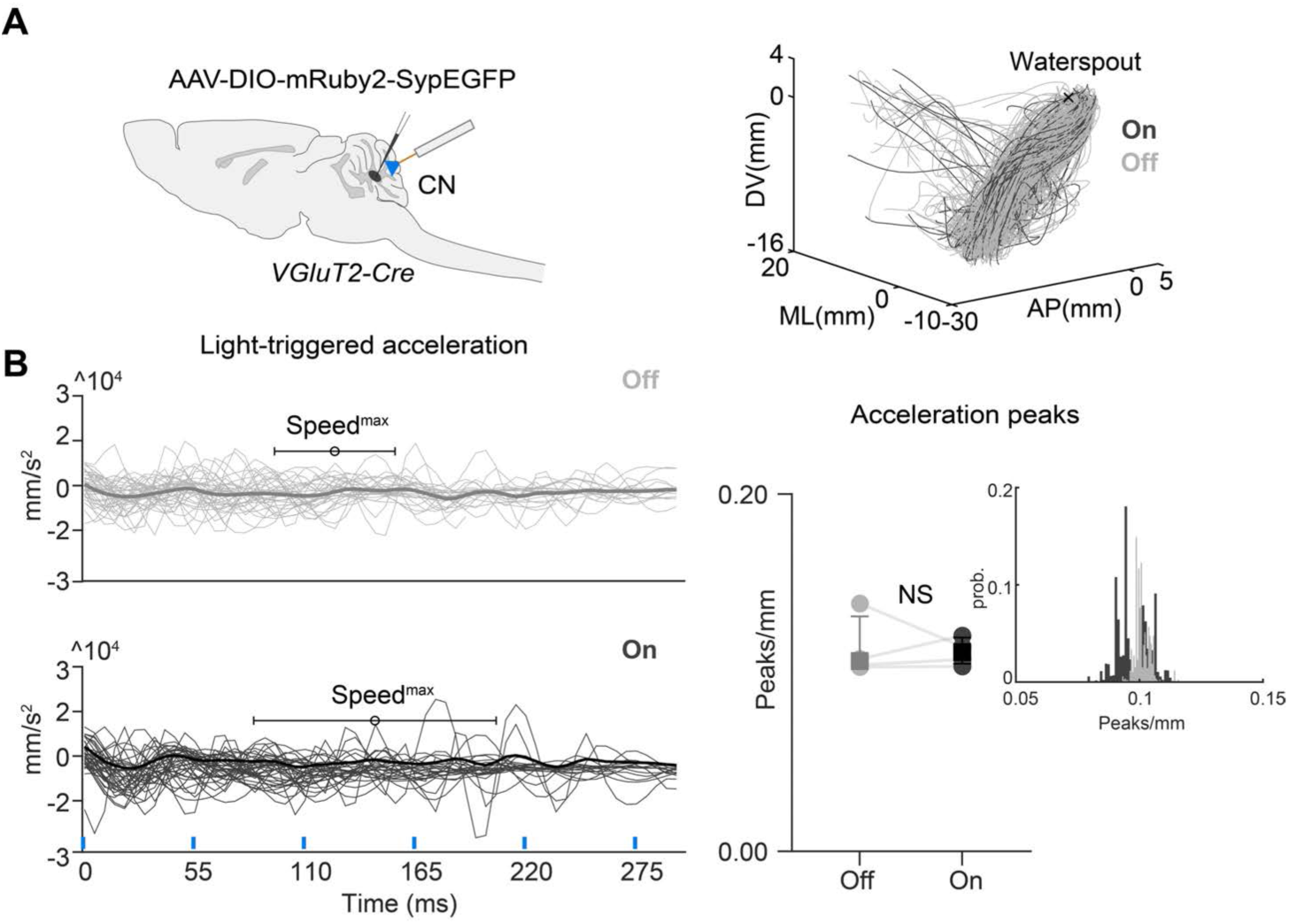
Photostimulation in control animals does not evoke directional limb movements. **(A)** Strategy for viral expression of fluorophores in excitatory CN neurons (left). 3D trajectories of all trials from an example animal; grey lines indicate light off trials, black lines indicate light on trials (right). **(B)** Hand acceleration triggered at light onset (black) compared to a similar timepoint during light off trials (grey) from a representative animal; thicker lines represent mean, thinner lines represent single trials, blue lines indicate pulses of light, distributions of when Speed^max^ occurred are shown (mean ± SEM) (left). Number of acceleration peaks normalized to path length across animals (right; N=4; NS, p=0.8203, paired t-test); circles represent averages within animal, squares and error bars represent median and IQR across mice. Hierarchical bootstrapped distribution of medians (inset).

**fig. S6.**
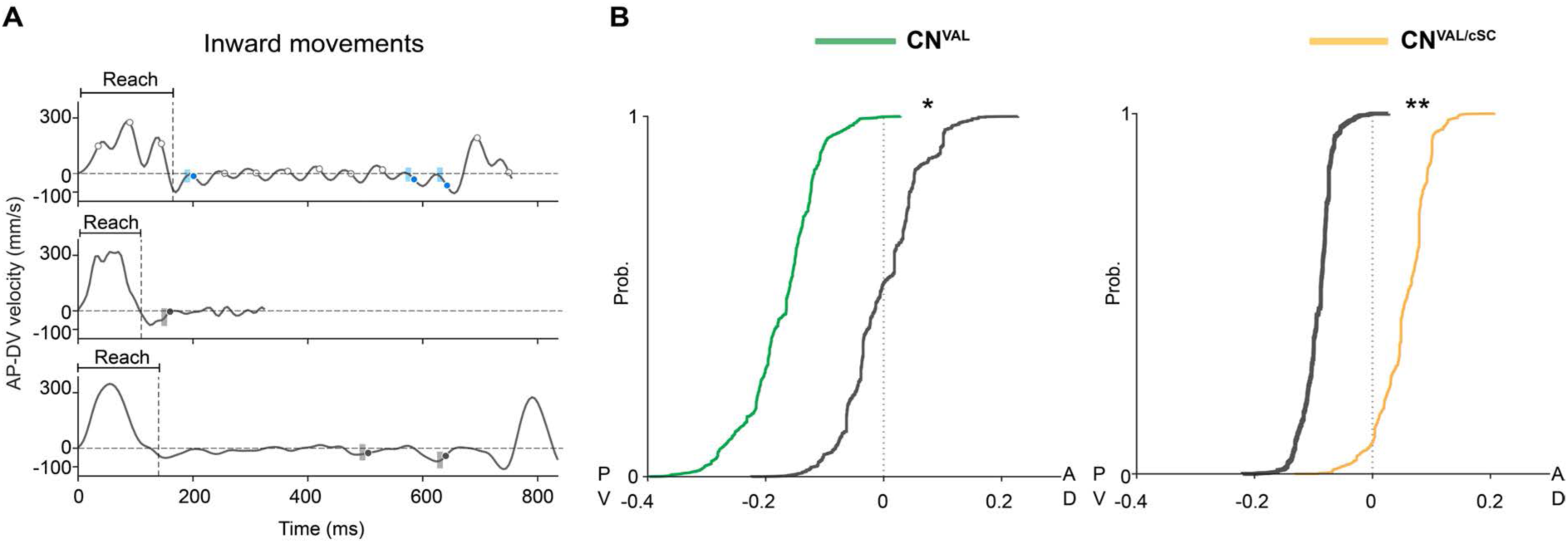
CN networks drive limb movements in opposing directions during inward movement. **(A)** Strategy for identifying stimulation timepoints during brief phases of the movement when the hand moves inward in AP-DV space. Blue circles in the top trace indicate light pulses during inward phases of the movement. White circles in the top trace indicate light pulses when the hand is not moving inward. Grey circles in the middle and bottom panel represent timepoints in unstimulated trials matched to the light on events by state (Materials and Methods). To be classified as an inward phase, the hand must be moving inward for at least 15 ms (blue and grey shaded regions). **(B)** AP-DV component of the stimulus-triggered submovements (green) compared to matched light off timepoints (black) across all CN^VAL^ animals (hierarchical bootstrapped CDF of medians; N=9; P-boot=0.023, Off>On, by joint probability) (left). AP-DV component of the stimulus-triggered submovements (yellow) compared to matched light off timepoints (black) across all CN^VAL/cSC^ animals (hierarchical bootstrapped CDF of medians; N=10; P-boot=0.004, On>Off, by joint probability) (right).

**fig. S7.**
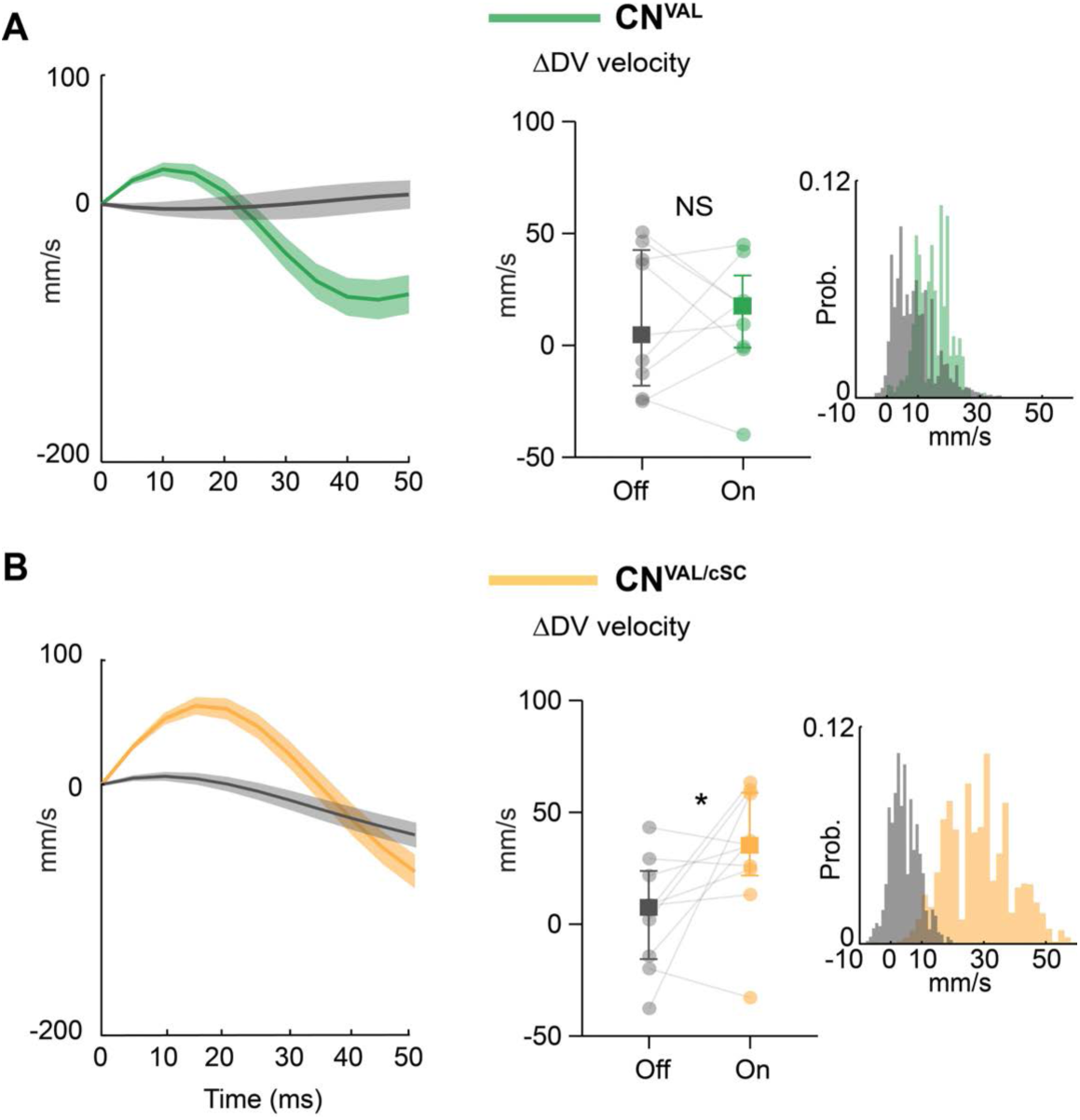
Differential effects on dorso-ventral (DV) movements caused by CN^VAL^ versus CN^VAL/cSC^ neuronal activation. **(A)** Change in DV velocity after light onset during light on trials (green) compared to a similar timepoint during light off trials (grey) from an example CN^VAL^ mouse. Lines indicate mean, shaded regions indicate SEM (left). No change in DV velocity in a 5 ms window 20–25 ms after light onset during light on trials compared to a similar timepoint during light off trials across CN^VAL^ mice (right, light off; median=4.535 mm/s, IQR=60.730, light on; median=17.430 mm/s, IQR=32.141; N=9; NS, p=0.989, paired t-test). Circles represent averages within animal, squares and error bars represent median and IQR across mice. Hierarchical bootstrapped distribution of medians (inset). **(B)** Change in DV velocity after light onset during light on trials (yellow) compared to a similar timepoint during light off trials (grey) from an example CN^VAL/cSC^ mouse. Change in DV velocity in a 5 ms window 20–25 ms after light onset during light on trials compared to a similar timepoint during light off trials across CN^VAL/cSC^ mice (right, light off; median=7.459 mm/s, IQR=39.300, light on; median=35.240 mm/s, IQR=37.010; N=10; p=0.041, paired t-test). Hierarchical bootstrapped distribution of medians (inset).

**fig. S8.**
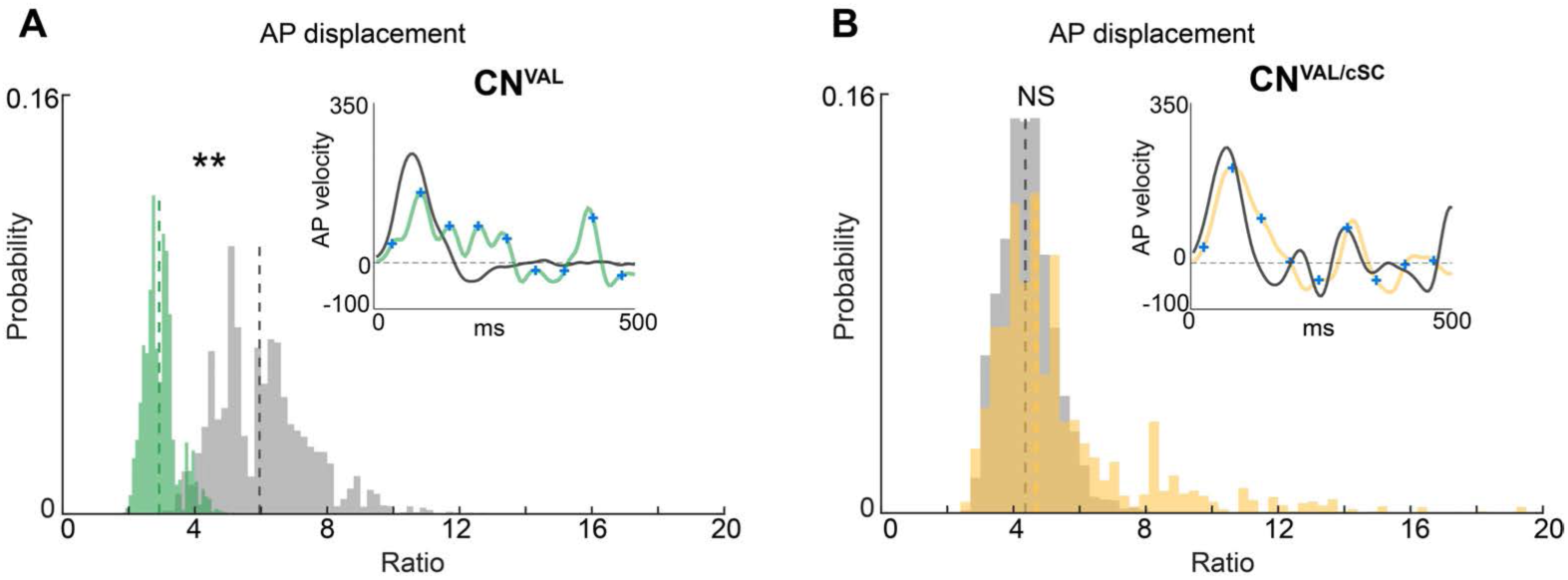
Differential effects on forward limb displacement throughout the reach when activating CN^VAL^ and CN^VAL/cSC^ neurons. **(A)** Total forward displacement during the reach is reduced when CN^VAL^ neurons are photoactivated; histograms show hierarchical bootstrapped distribution of medians, dotted lines show the median across mice (N=9; P-boot=0.0054, Off>On, by joint probability). Inset shows an example AP velocity trace, light off (grey), light on (green), from which AP displacement (the ratio of forward displacement / backward displacement) is calculated. Blue crosses represent light onset. **(B)** Same as (A), for photoactivation of CN^VAL/cSC^ neurons. Forward displacement in the light off and light on conditions is similar, due to the fact that movement throughout the reach is naturally in the outward direction; histograms show hierarchical bootstrapped distribution of medians, dotted lines show the median across mice (N=10; NS, P-boot=0.576).

**fig. S9.**
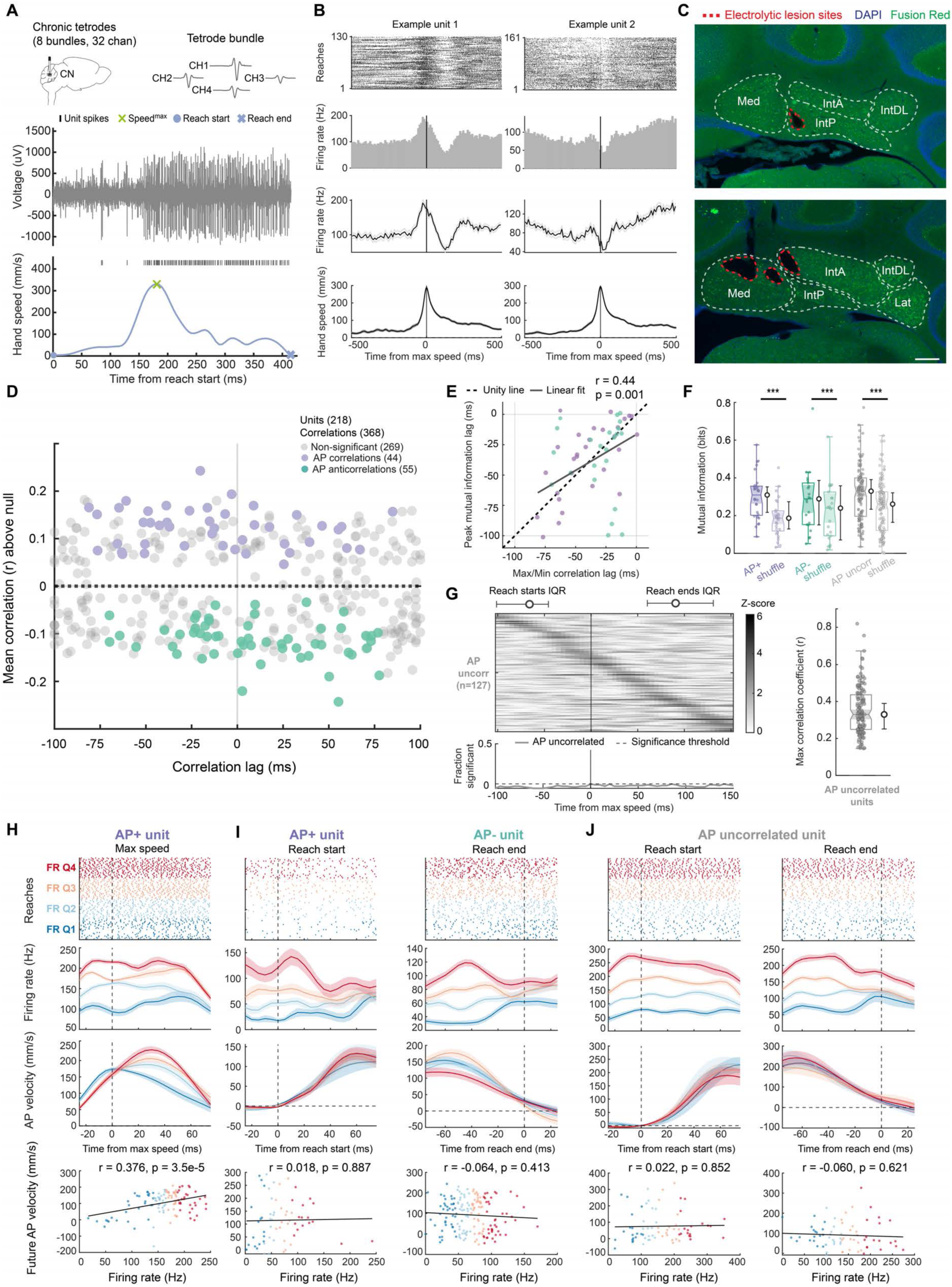
Extracellular recordings of CN neurons. **(A)** Schematic of chronic tetrode recordings in CN (top left) and example unit waveforms recorded across all channels of a tetrode bundle (top right). Example voltage traces, unit spikes, and hand speed across a single reach (bottom). **(B)** Two example unit rasters (top), binned firing rate (top middle), binned and smoothed firing rate (bottom middle; lines indicate mean, shaded regions indicate SEM), and hand speed (bottom). **(C)** Sections from two recorded animals showing electrolytic lesion sites from tetrodes. Scale bar: 250 μm. **(D)** All significant positive (N=44, purple) and negative (N=55, teal) correlations of firing rate to AP velocity above the trial shuffled null across all recorded units (N=6 mice; 218 units). The total number of significant and non-significant correlations exceeds unit number because many units show one peak at negative lag and another peak at positive lag with opposite sign. Positive lag peaks might reflect sensory responses and were excluded from subsequent analysis. **(E)** Correlation lag and peak mutual information lag of AP+ (purple) and AP-(teal) units, fit with a linear model (r and p values shown). **(F)** Mutual information between FR and AP velocity of AP+, AP-, and AP uncorrelated units at peak mutual information lag compared to trial shuffled controls (left to right: N=5 animals, 28 units, P-boot=3.0e-4; N=5 animals, 22 units, P-boot=8.0e-4; N=6 animals, 127 units, P-boot=3.0e-4, hierarchical bootstrap comparison with Holm-Bonferroni correction for multiple comparisons). **(G)** Sliding window z-scored correlations (absolute value) between firing rate and future AP velocity relative to maximum hand speed for all AP uncorrelated units (top, N=6 mice, 127 units), with median and IQR of reach starts and ends (top). Fraction of significantly correlated AP uncorrelated units over time, relative to maximum hand speed (bottom). Absolute value of maximum correlation coefficients for AP uncorrelated units with hierarchical bootstrapped median values and 95% confidence interval (right). Sorting uncorrelated units by peak in a time varying signal is expected to produce a uniform distribution across the time window, as seen. Note the lack of units crossing significance threshold (as compared to Fig. 3C). **(H)** Example AP+ unit at another reach phase (also see Fig. 3D). Raster across all reaches binned into firing rate (FR) quartiles (defined by 0 to +50 ms after time zero, top), smoothed firing rate (middle top), and AP velocity (middle bottom), relative to maximum speed. Bottom plot shows unit correlation of FR with future AP velocity, determined by unit’s cross-correlation lag, fit with a linear model (r and p values shown). **(I)** Example reach phases when AP+ and AP-units show no correlation with future AP velocity. Same as (H) for an example AP+ unit (left) and AP-unit (right) aligned to reach start (left) and reach end (right). **(J)** Same as (H) for an example AP uncorrelated unit showing no correlation with future AP velocity aligned to reach start (left) and reach end (right).

**fig. S10.**
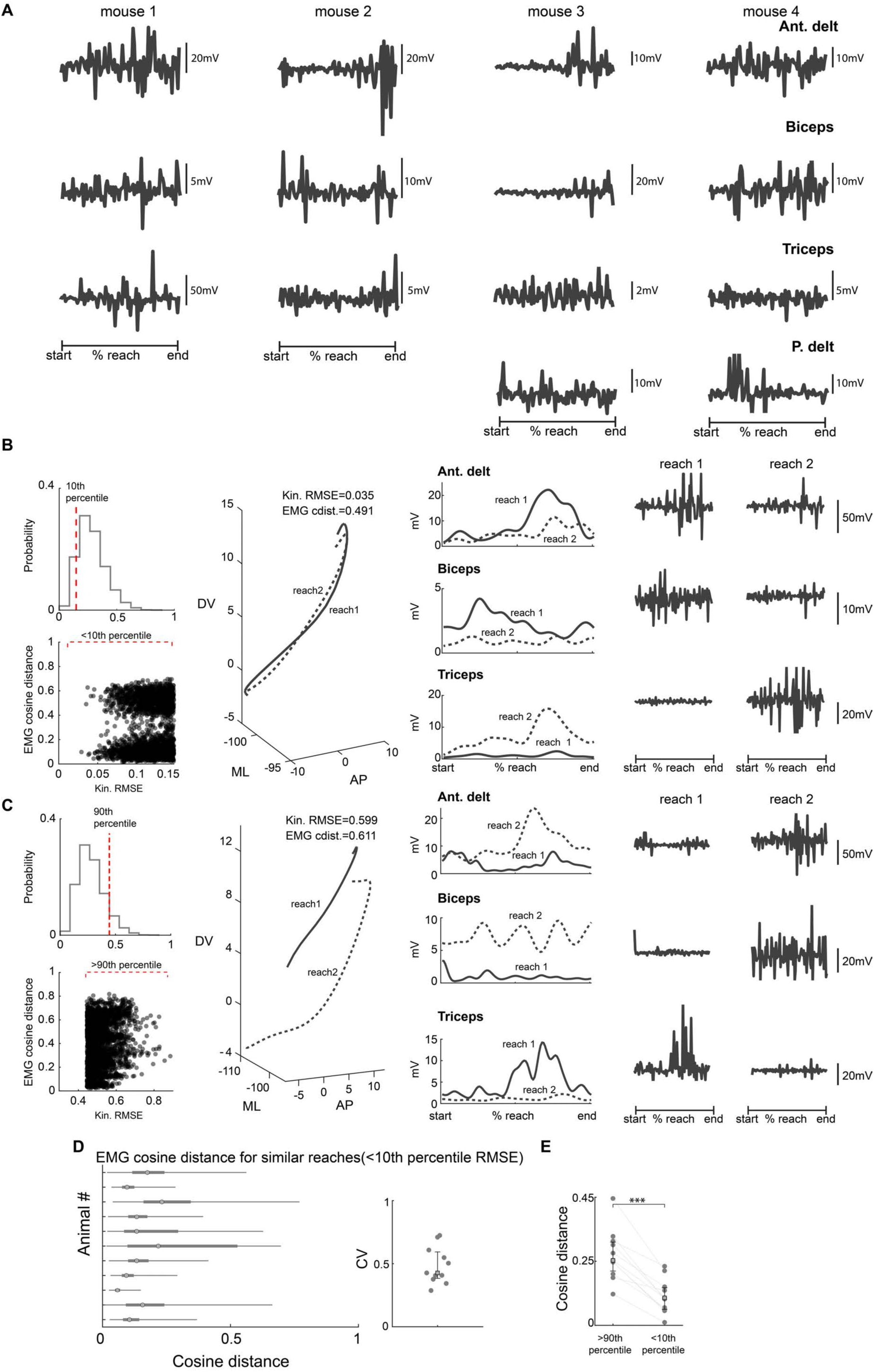
Variability in muscle recruitment across and within mice during reaching. **(A)** Example EMG traces showing variability in muscle recruitment during time-normalized reaches between animals. **(B)** Distribution of RMSE values of pairwise kinematic (kin) comparisons in an example animal during unperturbed reaches. Dotted red line indicates the most similar 10^th^ percentile cut-off value (top, left). EMG cosine distance (cdist) between reach pairs with RMSE values < 10^th^ percentile cut off (bottom, left). Example reach pair with low kinematic RMSE; reach 1, solid line; reach 2, dotted line (middle, left); scales in mm. EMG envelope for the two reaches (middle, right). Raw EMG traces for same reaches (right). **(C)** Distribution of RMSE values of pairwise kinematic (kin) comparisons in an example animal during unperturbed reaches. Dotted red line indicates the least similar 90^th^ percentile cut-off value (top, left). EMG cosine distance (cdist) between reach pairs with RMSE values > 90^th^ percentile cut off (bottom, left). Example reach pair with high kinematic RMSE; reach 1, solid line; reach 2, dotted line (middle, left); scales in mm. EMG envelope for the two reaches (middle, right). Raw EMG traces for same reaches (right). **(D)** Range of EMG cosine distance values for the most similar reach pairs (RMSE<10^th^ percentile cut-off). Each row is one animal, circle is median, thin line is range, thicker bar is IQR (left). Coefficient of variation (CV) of cosine distance values across animals (right) (N=11). **(E)** Cosine distance values compared between low-similarity reach pairs (RMSE>90th percentile cut-off) and high-similarity reach pairs (RMSE<10^th^ percentile cut-off) (N=11 mice; p=0.00001, paired t-test).

**fig. S11.**
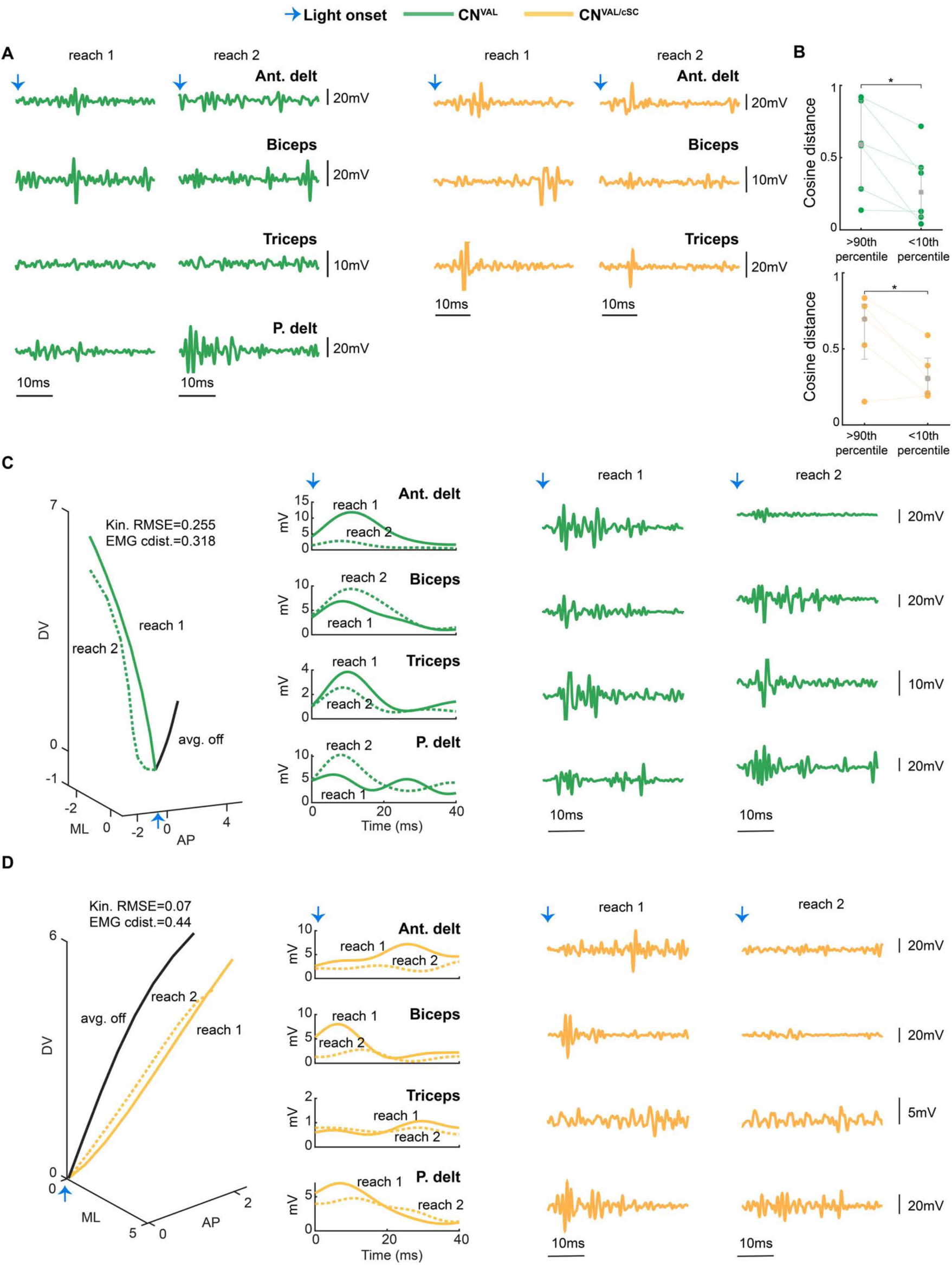
Variability in muscle recruitment after CN^VAL^ and CN^VAL/cSC^ stimulation. **(A)** Raw EMG traces corresponding to the EMG envelope traces shown in Fig. 4B,**C**; CN^VAL^ (left), CN^VAL/cSC^ (right). **(B)** Cosine distance values compared between low-similarity reach pairs (RMSE>90th percentile cut-off) and high-similarity reach pairs (RMSE<10^th^ percentile cut-off). CN^VAL^ (top; N=6; p=0.0247, paired t-test); CN^VAL/cSC^ (bottom; N=5; p=0.036, paired t-test). **(C)** Additional example of a reach pair from a CN^VAL^ animal with kinematic RMSE<10^th^ percentile cut-off; reach 1, solid line; reach 2, dotted line; average light off trajectory (black) shown for reference (left); scales in mm. EMG envelope (middle) and raw EMG traces for the two reaches (right). **(D)** Same as (C) for a CN^VAL/cSC^ animal. Blue arrows indicate light onset.

**fig. S12.**
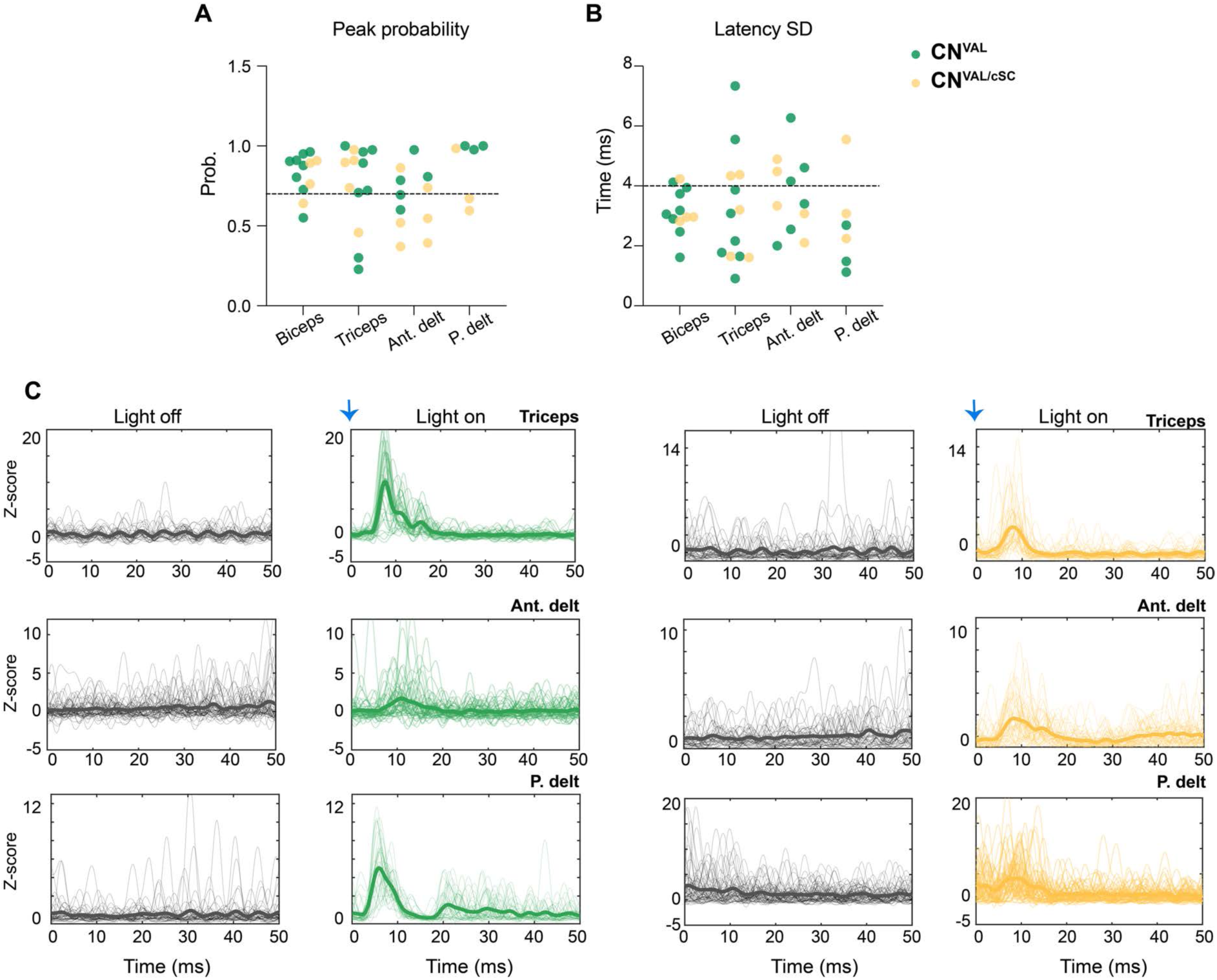
Thresholds for determining latency of muscle recruitment during CN^VAL^ and CN^VAL/cSC^ neuronal stimulation. **(A)** Probability of detecting a peak in z-scored muscle activity one standard deviation above the mean and less than 20 ms after light onset; dotted line indicates 70% threshold. **(B)** Standard deviation (SD) of the latency of the peak in EMG activity after light onset; dotted line indicates threshold SD of 4 ms (see **table S2**). **(C)** Light-triggered averaging of z-scored EMGs from example animals in light off (left) and light on (right) conditions. Green, example CN^VAL^ mice; yellow, example CN^VAL/cSC^ mice. Thicker lines represent mean; thinner lines represent single trials. Light onset (blue arrow) aligned to zero in light on plots.

**table S1.** Regions receiving significant projections from CN^VAL^ and CN^VAL/cSC^ pathways. Nomenclature and abbreviations are listed according to the Common Coordinate Framework v3.0, Allen Institute for Brain Science. Mean ± SEM for percentage distribution of pixels and density and are listed for each identified region (Materials and Methods).

| **Abbreviation** | **Region** | **Laterality** | **CN<sup>VAL</sup>**<br>distribution<br>(%, mean±SEM) | **CN<sup>VAL/cSC</sup>**<br>distribution<br>(%, mean±SEM) | **CN<sup>VAL</sup>**<br>density<br>(%, mean±SEM) | **CN<sup>VAL/cSC</sup>**<br>density<br>(%, mean±SEM) |
|---|---|---|---|---|---|---|
| VAL | Ventral anterior lateral complex of the thalamus | Contralateral | 2.333 ± 0.251 | 4.278 ± 1.498 | 3.017 ± 1.493 | 0.814 ± 0.172 |
| VM | Ventral medial nucleus of the thalamus | Contralateral | 1.630 ± 0.266 | 1.708 ± 0.687 | 2.259 ± 1.135 | 0.374 ± 0.117 |
| ZI | Zona Incerta | Contralateral | 2.665 ± 0.740 | 0.819 ± 0.266 | 2.403 ± 1.396 | 0.089 ± 0.025 |
| FF | Fields of forel | Contralateral | 0.498 ± 0.209 | 0.764 ± 0.175 | 3.029 ± 1.847 | 0.568 ± 0.268 |
| APN | Anterior pretectal nucleus | Contralateral | 2.802 ± 0.562 | 0.944 ± 0.407 | 2.219 ± 1.250 | 0.113 ± 0.028 |
| RN | Red nucleus | Contralateral | 4.431 ± 0.711 | 10.155 ± 2.362 | 4.347 ± 2.059 | 1.798 ± 0.455 |
| MRN | Midbrain reticular nucleus | Contralateral | 7.658 ± 1.578 | 10.644 ± 1.235 | 1.615 ± 0.791 | 0.375 ± 0.114 |
| SuCiw | Superior colliculus, motor related, intermediate white layer | Contralateral | 4.358 ± 0.210 | 0.299 ± 0.105 | 2.122 ± 1.042 | 0.029 ± 0.014 |
| SuCig | Superior colliculus, motor related, intermediate gray layer | Contralateral | 3.054 ± 0.529 | 0.397 ± 0.180 | 0.529 ± 0.970 | 0.035 ± 0.018 |
| TRN | Tegmental reticular nucleus | Contralateral | 1.300 ± 0.429 | 1.809 ± 0.425 | 2.799 ± 1.249 | 0.449 ± 0.181 |
| PRNc | Pontine reticular nucleus, caudal part | Contralateral | 1.668 ± 0.304 | 6.287 ± 2.222 | 0.799 ± 0.427 | 0.443 ± 0.170 |
| MARN | Magnocellular reticular nucleus | Contralateral | 0.852 ± 0.327 | 3.148 ± 0.668 | 2.121 ± 1.182 | 1.316 ± 0.581 |
| GRN | Gigantocellular reticular nucleus | Contralateral | 2.496 ± 1.227 | 11.766 ± 1.431 | 1.362 ± 0.774 | 0.883 ± 0.324 |
| PGRNl | Paragigantocellular reticular nucleus, lateral part | Contralateral | 0.459 ± 0.191 | 2.100 ± 0.359 | 0.754 ± 0.427 | 0.538 ± 0.231 |
| LRNm | Lateral reticular nucleus, magnocellular part | Contralateral | 0.286 ± 0.116 | 2.720 ± 0.575 | 0.651 ± 0.374 | 1.287 ± 0.722 |
| MDRNv | Medullary reticular nucleus, ventral part | Contralateral | 0.524 ± 0.265 | 1.767 ± 0.204 | 1.096 ± 0.629 | 0.568 ± 0.204 |
| PB | Parabrachial nucleus | Ipsilateral | 5.479 ± 3.128 | 1.677 ± 0.138 | 2.126 ± 0.634 | 0.307 ± 0.104 |
| SUV | Superior vestibular nucleus | Ipsilateral | 3.262 ± 1.385 | 0.615 ± 0.138 | 3.363 ± 0.685 | 0.321 ± 0.139 |
| PARN | Parvicellular reticular nucleus | Ipsilateral | 8.362 ± 2.971 | 1.120 ± 0.380 | 2.998 ± 1.424 | 0.100 ± 0.054 |
| MDRNd | Medullary reticular nucleus, dorsal part | Ipsilateral | 2.715 ± 0.429 | 0.264 ± 0.108 | 2.641 ± 1.229 | 0.066 ± 0.030 |

**table S2.** Number of mice with light-triggered muscle activity in the biceps brachii, triceps brachii, anterior deltoid, and posterior deltoid. Numerator denotes the number of animals with detected light-triggered muscle activity, and the denominator denotes the number of animals recorded for a given muscle within each group. See Materials and Methods for criteria.

| **Group** | **Biceps brachii** | **Triceps brachii** | **Anterior deltoid** | **Posterior deltoid** |
|---|---|---|---|---|
| CN<sup>VAL</sup> | N=7/8 | N=6/8 | N=3/5 | N=3/3 |
| CN<sup>VAL/cSC</sup> | N=3/4 | N=4/5 | N=2/6 | N=1/3 |

